# A benchmark of structural variation detection by long reads through a realistic simulated model

**DOI:** 10.1101/2020.12.25.424397

**Authors:** Nicolas Dierckxsens, Tong Li, Joris R. Vermeesch, Zhi Xie

## Abstract

Despite the rapid evolution of new sequencing technologies, structural variation detection remains poorly ascertained. The high discrepancy between the results of structural variant analysis programs makes it difficult to assess their performance on real datasets. Accurate simulations of structural variation distributions and sequencing data of the human genome are crucial for the development and benchmarking of new tools. In order to gain a better insight into the detection of structural variation with long sequencing reads, we created a realistic simulated model to thoroughly compare SV detection methods and the impact of the chosen sequencing technology and sequencing depth. To achieve this, we developed Sim-it, a straightforward tool for the simulation of both structural variation and long-read data. These simulations from Sim-it revealed the strengths and weaknesses for current available structural variation callers and long read sequencing platforms. Our findings were also supported by the latest structural variation benchmark set developed by the GIAB Consortium. With these findings, we developed a new method (combiSV) that can combine the results from five different SV callers into a superior call set with increased recall and precision. Both Sim-it and combiSV are open source and can be downloaded at https://github.com/ndierckx/.

## INTRODUCTION

In order to decipher the genetic basis of human disease, a comprehensive knowledge of all genetic variation between human genomes is needed. Until recently, the emphasis has been on single-nucleotide polymorphisms, as these variants are easier to trace with current sequencing technologies and algorithms (1, 2). Over the past 20 years, we gained a better view on the prevalence of structural variation (SV), which changed our perspective on the impact it has on genomic disorders. We now know that structural variation contributes more to inter-individual genetic variation at the nucleotide level than single nucleotide polymorphisms (SNPs) and short indels together (3, 4). Structural variation covers insertions, deletions, inversions, duplications and translocations that are at least 50 bp in size. The limited length of Next-Generation Sequencing (NGS) reads (≤ 300 bp) hampers the detection of SVs, especially for insertions (3, 5). These technical limitations can be partially overcome by the third-generation sequencing, which is capable of producing far longer read lengths (6, 7). The race for dominance on the third-generation sequencing market has significantly reduced the costs per Mb and increased the throughput and accuracy, which makes these technologies (PacBio and Oxford Nanopore) currently the best option for structural variance detection (8).

The downside of these longer reads are their lower accuracies (85-95%) compared to NGS reads (> 99%), which requires new computational tools to achieve an optimal SV detection. Even though several algorithms were developed over the past decade, there is a large discrepancy between their outputs. Assessing the performance of SV detection tools is not straightforward, as there is no gold standard method to accurately identify structural variation in the human genome. To overcome this shortcoming, the Genome in a Bottle (GIAB) Consortium recently published a sequence-resolved benchmark set for identification of SVs, though it only includes deletions and insertions not located in segmental duplications (9). For as long as there is no completely resolved benchmark available, it is crucial to simulate a human genome with a set of structural variations that resembles reality as close as possible. There are a wide range of structural variation and long sequencing reads simulators available, yet without a thorough benchmark, it is impossible to know which tools are best suited to design the model you want to simulate. Therefore we compared several structural variance and long read simulators for their system requirements and available features. Furthermore we introduce Sim-it, a new SV and long read simulator that we designed for the assessment of SV detection with long read technologies.

The most complete structural variance detection study to date identified around 25,000 SVs for each individual by combining a wide range of sequencing platforms (3). The large amount of sequencing data used for this study makes it too costly to reproduce it on a larger scale, but it can be used as a golden standard for other SV studies. We used the results of this study to produce a realistic model for the evaluation of the available SV detection algorithms and to develop a new script that can improve SV detection by combining the results of existing tools.

## RESULTS

### Structural variation simulation benchmark

We compared the features and computational resources of five structural variation simulators, as shown in Table 1. Although all simulators can simulate the most common types of structural variation (insertions, deletions, duplications, inversions and translocations), more complex SV events need to be included in order to reproduce a realistic SV detection model. For Sim-it, we also included complex substitutions and inverted duplications, both common types of variation in germline and somatic genomes (5, 10, 11, 12). Additionally, it is possible to combine random generated SV events with a defined list of SVs at base pair resolution. Random generated SVs will be distributed realistically across the genome with higher prevalence around the telomeres. As output, Sim-it produces a sequence file in FASTA format and optionally long sequencing reads (PacBio or ONT). Although none of the other tools has a proprietary method to simulate long reads, Varsim can generate long reads through PBSIM or LongISLND. Currently, Sim-it does not support short read or phylogenetic clonal structure simulation. As for computational resources, Sim-it performed best on peak memory consumption and runtime. With 1 GB as peak memory consumption and 5 min 30 s as runtime (single core) to simulate 24,600 SV events, Sim-it can be implemented for any set of SVs on a small desktop or laptop. SVEngine and Varsim also have relatively low runtimes, though a peak memory consumption of respectively 24.3 GB and 8 GB limits it’s use on machines with limited computational resources. SCNVsim was excluded as it does not accept a set list of SVs as input and has an upper limit of 600 SVs for random simulation.

**Table 1.** Available features and system requirements of structural variation simulators.

|  | Sim-it | SVEngine | RSVSim | Varsim | SCNVsim |
| --- | --- | --- | --- | --- | --- |
| **INPUT** |  |  |  |  |  |
| INS, DEL, INV, DUP and TRA | ✓ | ✓ | ✓ | ✓ | ✓ |
| Inverted duplications | ✓ | ✓ |  | ✓ |  |
| Complex substitutions | ✓ |  | ✓ | ✓ |  |
| Foreign sequence insertion | ✓ | ✓ |  | ✓ |  |
| Random generated SVs | ✓ | ✓ | ✓ |  | ✓ |
| Realistic distribution of random SVs | ✓ |  | ✓ |  |  |
| Breakpoint at base pair resolution | ✓ | ✓ | ✓ | ✓ |  |
| **OUTPUT** |  |  |  |  |  |
| Separate haplotypes | ✓ | ✓ |  | ✓ | ✓ |
| Short sequencing reads |  | ✓ |  | ✓ |  |
| Long sequencing reads | ✓ |  |  | ✓ |  |
| Graphical output | ✓ |  | ✓ |  |  |
| Phylogenetic clonal structure |  | ✓ |  |  | ✓ |
| **COMPUTATIONAL RESOURCES** |  |  |  |  |  |
| Wall time | 5 m 30 s | 12 m 04 s | 938 m | 9 m 27 s | / (*) |
| Virtual Memory | 1 GB | 24.3 GB | 11.9 GB | 8 GB | / (*) |

\*SCNVsim was excluded from the benchmark.

### Long read simulation benchmark

We assessed the quality of the simulated long reads by comparing their error profiles to those of real PacBio and ONT sequencing reads. Additionally, we compared the features and system requirements for each tool.

Several systems of ONT and PacBio technologies have been released in the last decade, each with different specifications for the sequencing reads. This complicates an accurate simulation as a specific error profile is needed for each released system. From the 8 tested simulators, only Sim-it, Badread and LongISLND support simulations for both ONT and PacBio. Sim-it provides error profiles for ONT, PacBio RS II, PacBio Sequel II and Pacbio Sequel HiFi systems, while other simulators are limited to one or two error profiles. This shortcoming can be overcome by training a new model for a system, a feature supported by all simulators apart from PBSIM and SimLoRD. This is more laborious and a real dataset along with an accurate reference sequence is required to train a new model. Not all updates require a completely new error profile, therefore we provide the option to adjust the overall accuracy and read length independently from the error profile. As for computational resources, PBSIM performed the best with just 5 minutes and 0.25 GB of RAM to simulate 15x coverage for chromosome 1 of GRCh38. Besides for DeepSimulator, Badread and NanoSim, computational resources stayed within a reasonable range.

Available features and computational resources determine the suitability and user-friendliness of the simulators, but not the accuracy of the simulation. Therefore, we compared the context-specific error patterns of the simulated reads to real long sequencing datasets. Figure 1A shows the context-specific errors derived from real data from Nanopore PromethION and PacBio Sequel II sequencing reads, as well from their respective simulations by Sim-it. These context-specific error heat-maps were generated for each of the 8 simulators and can be found in Supplementary materials. NanoSim generated random errors in stead of a context-specific error pattern, while PBSIM and SimLoRD have simplified patterns. For Sim-it, the length of deletions and insertions closely match the real data (Figure 1C and 1D). LongISLND has proportionally too many single nucleotide deletions, while the asymmetry for DeepSimulator is caused by a low absolute number of deletions, which is not adjustable.

**Figure 1.**
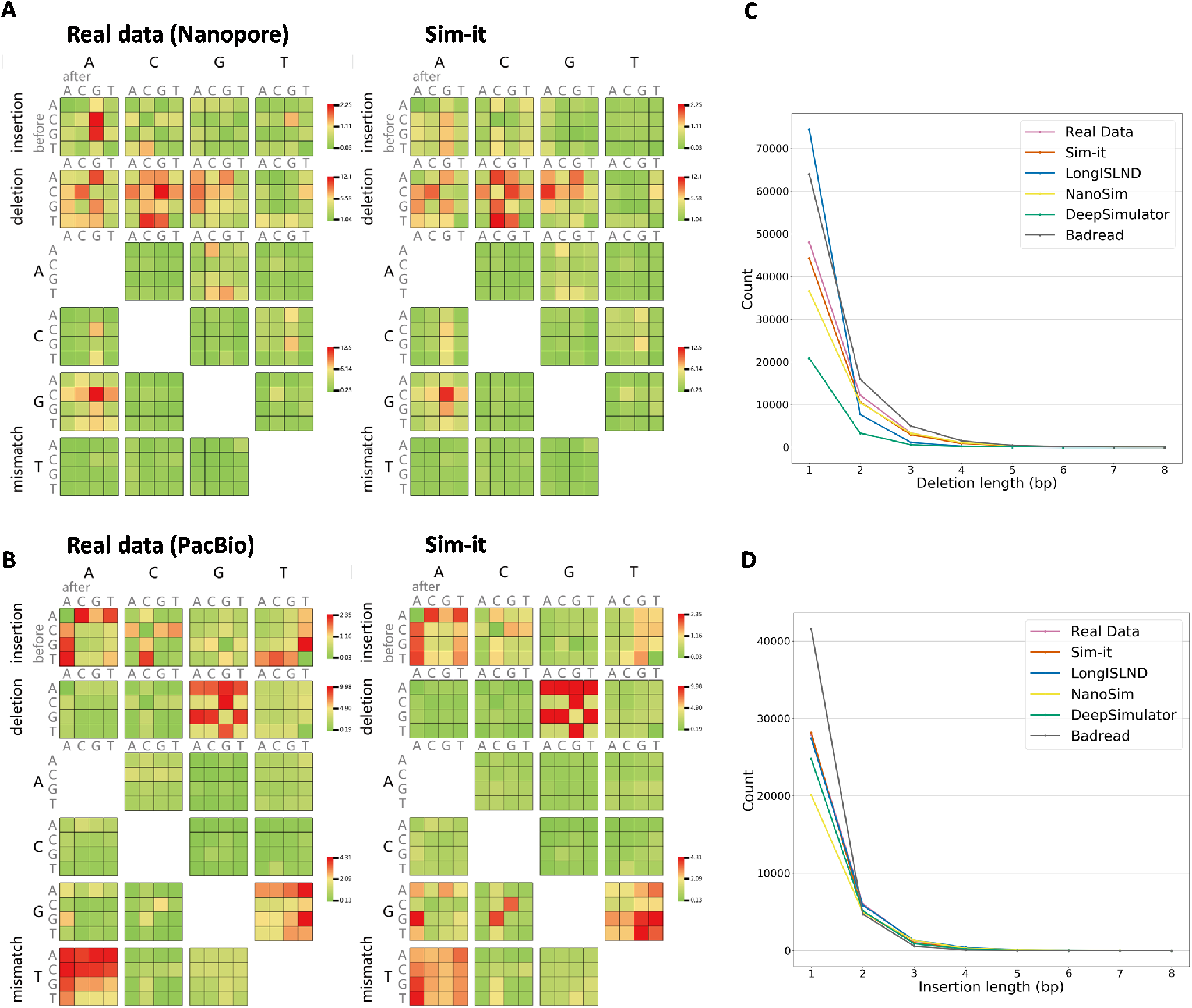
Context-specific error patterns for mismatches and indels. (A) Context-specific error patterns for real data of Nanopore (9.4.1) and simulated data from Sim-it. (B) Context-specific error patterns for real data of PacBio Sequel II and simulated data from Sim-it. (C) Deletion lengths for real Nanopore data and the simulations of the benchmarked tools. (D) Insertion lengths for real Nanopore data and the simulations of the benchmarked tools.

### Structural variance detection using simulated long reads

We assessed the performance of 6 long read SV detection algorithms through a realistic model of 24,600 SV events. Additionally, we made a comparison between PacBio and ONT technology and evaluated the impact of the read length and coverage depth. For each simulated dataset, a separate score for each type of SV and for the four essential parameters that define SVs; namely position, length, type and haplotype were calculated.

We performed a complete analysis on each of the 6 SV callers for a Nanopore and a PacBio Sequel II long reads and a HiFi reads dataset with a sequencing depth of 20x (Table 2). For each dataset, Picky had more than 19,000 false positives and false negatives, with an outlier of 46,502 false positives for the PacBio HiFi dataset. We therefore excluded Picky for any further analysis or graphical output. All the statistics of Picky for all three 20x coverage datasets can be examined in the Supplementary Data.

**Table 2.** Benchmark statistics on three simulated datasets of 24,600 SVs for 5 existing SV callers and combiSV (combiSV (3): pbsv, Sniffles and SVIM combined; combiSV (5): all 5 tools combined).

|  |  | Sniffles | pbsv | NanoVar | NanoSV | SVIM | combiSV (3) | combiSV (5) |
| --- | --- | --- | --- | --- | --- | --- | --- | --- |
| Nanopore | Recall | 79.89% | 73.19% | 80.34% | 83.19% | 80.95% | 81.24% | 81.11% |
|  | Precision | 96.93% | 94.61% | 86.06% | 87.56% | 91.90% | 97.10% | 97.72% |
|  | Perfect matches | 11.62% | 34.94% | 1.58% | 0.52% | 6.32% | 31.84% | 31.88% |
|  | Position score | 87.6% | 90.6% | 78.6% | 88.7% | 82.2% | 90.7% | 90.5% |
|  | Length score | 90.9% | 95.2% | 83.4% | 90.3% | 90.4% | 95.5% | 95.3% |
|  | Type score | 93.6% | 94.8% | 87.7% | 45.6% | 94.8% | 93.7% | 94.4% |
|  | Haplotype score | 58.4% | 94.6% | 88.6% | 94.9% | 93.0% | 94.8% | 94.7% |
|  | **Total score** | **64.3%** | **64.0%** | **54.0%** | **56.1%** | **64.5%** | **73.4%** | **73.6%** |
| PacBio | Recall | 78.66% | 73.69% | 80.51% | 83.30% | 80.99% | 80.27% | 80.81% |
|  | Precision | 97.45% | 94.53% | 85.95% | 88.17% | 92.48% | 97.70% | 98.19% |
|  | Perfect matches | 12.82% | 47.00% | 2.24% | 4.30% | 7.76% | 43.37% | 42.99% |
|  | Position score | 87.4% | 91.6% | 79.1% | 88.3% | 82.7% | 91.6% | 91.6% |
|  | Length score | 93.0% | 96.5% | 79.9% | 93.4% | 92.5% | 96.7% | 96.7% |
|  | Type score | 95.2% | 94.7% | 87.4% | 46.0% | 95.1% | 95.2% | 95.2% |
|  | Haplotype score | 58.5% | 94.7% | 88.2% | 95.1% | 92.8% | 94.4% | 94.4% |
|  | **Total score** | **64.2%** | **64.9%** | **53.5%** | **57.3%** | **65.6%** | **73.6%** | **73.6%** |
| PacBio HiFi | Recall | 70.90% | 73.04% | 82.19% | 80.27% | 76.83% | 76.71% | 77.11% |
|  | Precision | 97.88% | 96.16% | 86.04% | 94.84% | 95.81% | 97.70% | 98.10% |
|  | Perfect matches | 26.12% | 59.46% | 12.90% | 29.68% | 57.49% | 57.26% | 57.21% |
|  | Position score | 93.0% | 93.1% | 86.3% | 92.9% | 92.6% | 93.1% | 93.0% |
|  | Length score | 97.7% | 97.1% | 88.8% | 96.2% | 97.0% | 97.8% | 97.7% |
|  | Type score | 94.9% | 94.6% | 88.9% | 43.3% | 94.6% | 94.9% | 94.8% |
|  | Haplotype score | 47.2% | 93.2% | 91.5% | 96.6% | 95.0% | 91.4% | 93.5% |
|  | **Total score** | **58.8%** | **65.9%** | **59.3%** | **63.4%** | **69.1%** | **70.4%** | **71.3%** |
| GIAB (Nanopore) | Recall | 91.21% | 87.16% | 89.28% | 93.96% | 92.78% | 93.73% | 93.60% |
|  | Precision | 91.72% | 89.89% | 66.51% | 55.82% | 84.10% | 92.44% | 91.98% |
|  | Perfect matches | 0.09% | 28.16% | 0.68% | 0.53% | 2.36% | 26.16% | 25.90% |
|  | Position score | 67.7% | 75.6% | 61.3% | 73.0% | 62.4% | 73.7% | 73.6% |
|  | Length score | 80.4% | 87.0% | 62.3% | 77.4% | 79.8% | 86.2% | 86.3% |
|  | Type score | 94.7% | 97.5% | 61.0% | 48.0% | 95.0% | 97.1% | 97.3% |
|  | Haplotype score | 37.5% | 90.0% | 66.0% | 87.1% | 85.0% | 89.1% | 87.6% |
|  | **Total score** | **55.3%** | **64.4%** | **10.7%** | **0.0%** | **53.8%** | **71.1%** | **70.2%** |

For a sequencing depth of 20x, Sniffles and pbsv achieved the best overall performance across all sequencing platforms. Sniffles produced the lowest number of false positives independent from sequencing platform and coverage depth (Table2 and Figure2). For PacBio HiFi data, Sniffles performs significantly worse than pbsv, which can be explained by the shorter read lengths (Figure 3). Although pbsv generally has a lower recall, it calls SVs more accurately (position, length, type, haplotype) than any other tool, independent from the platform or coverage depth. Subsequently, this high accurateness results in a significant higher number of perfect matches compared to other tools. Perfect matches are SVs called with the correct type, haplotype, exact length and position. For PacBio CLR and PacBio HiFi reads, pbsv manages to call respectively 47% and 59.46% of the detected SVs perfectly, which is quite impressive compared to the other tools. Only SVIM achieved a similar percentage for PacBio HiFi reads (57.49%), however not for PacBio CLR reads (7.76%). The highest recall is achieved by NanoSV and to a certain extend NanoVar (only for PacBio HiFi), however this is at the expense of a disproportional amount of false positives.

**Figure 2.**
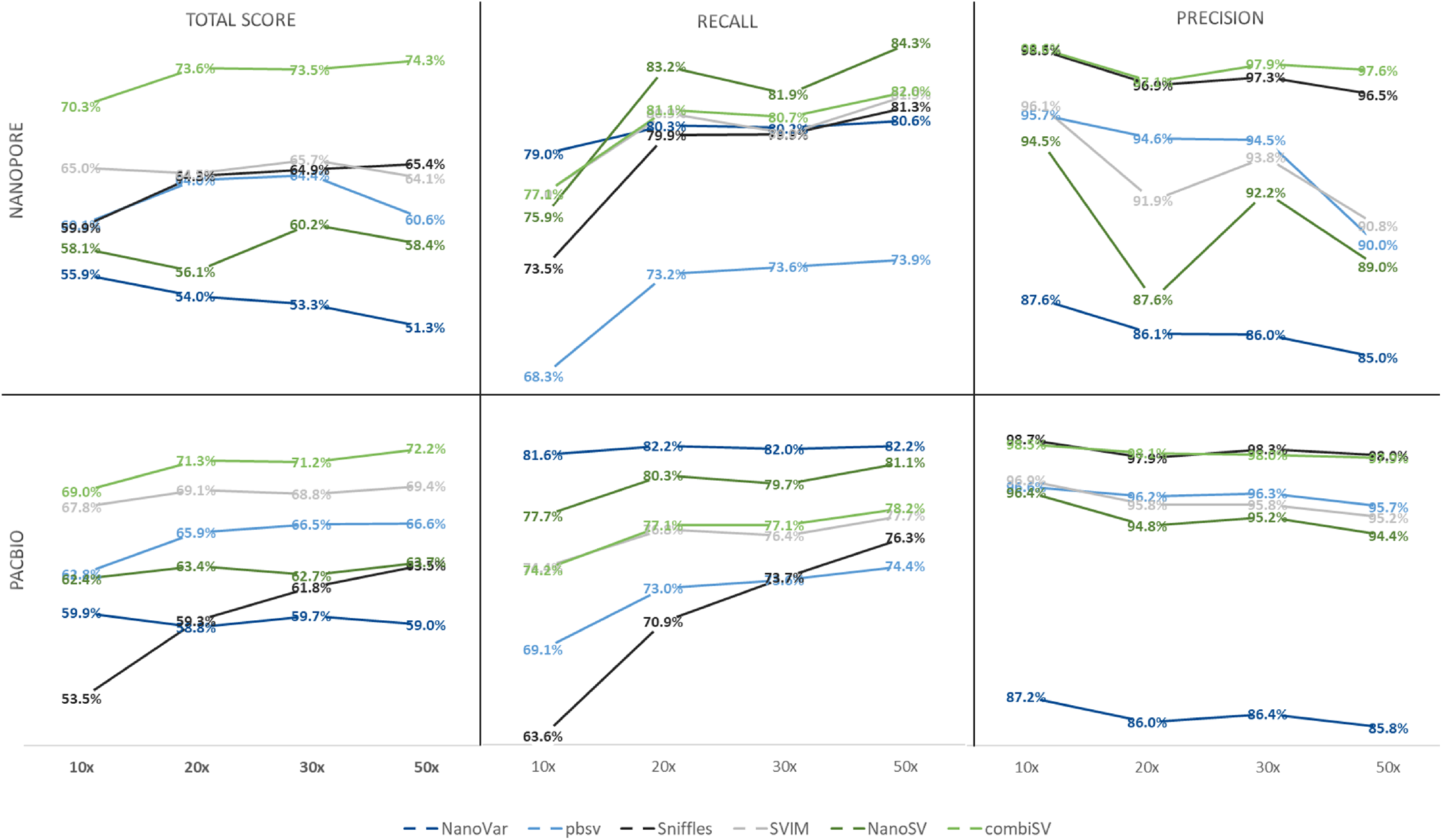
Structural variance detection stats for a series of Nanopore and PacBio HiFi datasets with increasing sequencing depths.

The 24,600 SVs can be classified by 5 different types, namely deletions, insertions, duplications, inversions and complex substitutions. We calculated the recall and precision metrics for each type of SV; Table 3 shows the results for the Nanopore 20x dataset, data metrics for the PacBio 20x and PacBio HiFi 20x datasets reveal similar patterns and can be examined in the Supplementary Data. NanoSV only classifies insertions, other SVs are indicated as breakend (BND). None of the SV callers classify complex substitutions in their output, which explains the missing precision values for this type. These complex substitutions seem to be the most problematic, as their recall values are very low for each of the tools. Recall and precision values of inversions are also far below the average for each of the tools. The low precision value for duplications detected by NanoVar can be explained by the fact that a significant fraction of the insertions are typed as a duplication.

**Table 3.**
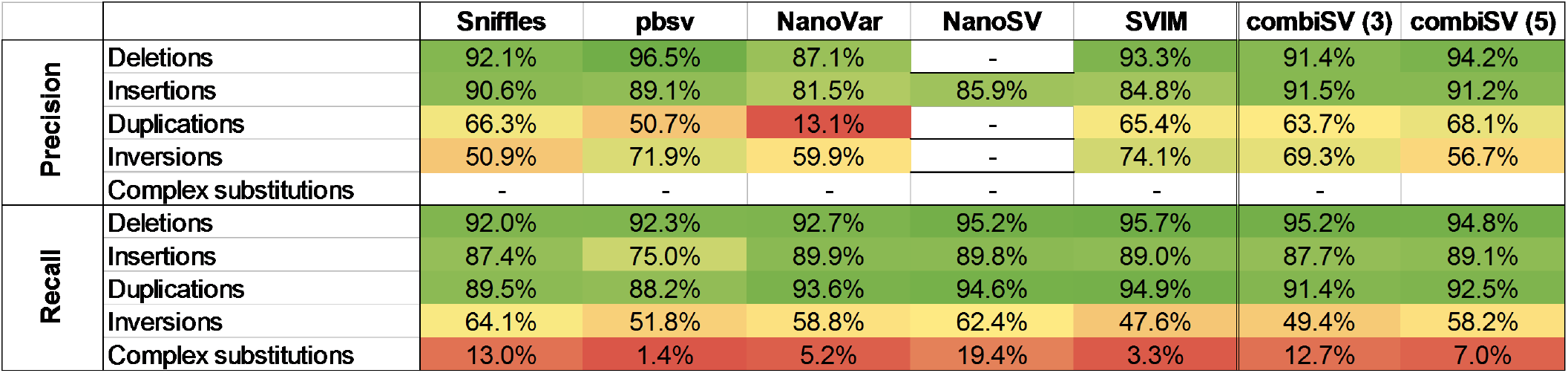
Precision and recall statistics for each type of SV from the Nanopore 20x dataset. (combiSV (3): pbsv, Sniffles and SVIM combined; combiSV (5): all 5 tools combined)

|  |  | Sniffles | pbsv | NanoVar | NanoSV | SVIM | combiSV (3) | combiSV (5) |
| --- | --- | --- | --- | --- | --- | --- | --- | --- |
| Precision | Deletions | 92.1% | 96.5% | 87.1% | - | 93.3% | 91.4% | 94.2% |
|  | Insertions | 90.6% | 89.1% | 81.5% | 85.9% | 84.8% | 91.5% | 91.2% |
|  | Duplications | 66.3% | 50.7% | 13.1% | - | 65.4% | 63.7% | 68.1% |
|  | Inversions | 50.9% | 71.9% | 59.9% | - | 74.1% | 69.3% | 56.7% |
|  | Complex substitutions | - | - | - | - | - | - | - |
| Recall | Deletions | 92.0% | 92.3% | 92.7% | 95.2% | 95.7% | 95.2% | 94.8% |
|  | Insertions | 87.4% | 75.0% | 89.9% | 89.8% | 89.0% | 87.7% | 89.1% |
|  | Duplications | 89.5% | 88.2% | 93.6% | 94.6% | 94.9% | 91.4% | 92.5% |
|  | Inversions | 64.1% | 51.8% | 58.8% | 62.4% | 47.6% | 49.4% | 58.2% |
|  | Complex substitutions | 13.0% | 1.4% | 5.2% | 19.4% | 3.3% | 12.7% | 7.0% |

To investigate the influence of increased sequencing coverage, we simulated 4 different datasets with sequencing depths of 10x, 20x, 30x and 50x for both Nanopore and PacBio HiFi (Figure 2). The general trends for increased sequencing depth are an increased recall and increased false positives, although depending on the tool, they can be disproportional to each other. NanoVar was designed to work on low sequencing depths and therefore does not display much gains in recall, yet a significant reduction in precision. Sniffles benefits the most from additional coverage with steep increases of recall together with a relatively low loss of precision. pbsv has a stable performance across all coverages, with the exception of Nanopore 50x, which exhibits a steep increase in false positives. The big drops in precision for NanoSV and SVIM at 20x and 50x coverage of Nanopore are caused by the additional filtering step we implemented for minimal variance allele coverage (3 for 10x and 20x, 5 for 30x and 50x). This shows how important the choice for the minimal coverage threshold is to obtain a good balance between recall and precision.

Besides sequencing depth, it is often believed that increasing sequencing lengths can improve assemblies and variance detection. We compared the SV detection metrics for three datasets of Nanopore with median read lengths of 15,000, 25,000 and 40,000 bp. We observed an increase in recall and overall score with increasing read lengths for each of the tools, with the most pronounced improvement from median lengths of 15k to 25k. NanoVar and pbsv show a modest rise in recall of 1% between 15k and 40k lengths, while Sniffles, SVIM, NanoSV and combiSV show an increase of 6%. All metrics of this comparison can be found in the Supplementary Data.

### Structural variance detection using real datasets

There is currently no SV call set covering the complete human genome that can be used as gold standard in a SV detection benchmark. The GIAB Consortium provides an accurate SV call-set of 5,260 insertions and 4,138 deletions, covering 2.5 GB of the human genome. Within the regions of the provided BED file, it is possible to accurately determine the recall and precision for both deletions and insertions. We benchmarked each of the tools for this high confidence set of SVs. We observed a similar pattern in benchmark metrics compared to the simulated dataset, with the exception of the low precision values for NanoVar and NanoSV. Recall values for the GIAB dataset are across all tools higher than for the simulated datasets, which can be explained by the exclusion of complex regions in the GIAB call set. The benchmark metrics of this real dataset also confirms our findings from the simulated datasets, though sometimes more outspoken in the real dataset. Sniffles has the highest precision, pbsv characterizes SVs the most accurate, NanoSV has the highest recall, low haplotype scores for sniffles and low position scores for SVIM are all findings that were observed with both simulated and real datasets.

We based our simulated datasets on a SV call set of NA19240 (nstd152), which was obtained through an elaborated SV study that combined a wide range of sequencing data (3). To compare our simulation to the original genome, we performed the same benchmark on a public available PacBio CLR dataset of that study. Recall and precision values of the real dataset were significantly lower, with an average of respectively 60% and 48%. An even more striking difference were the recall percentages of around 60% for complex substitutions, while these values ranged between 1% and 20% for the simulated datasets, independent from sequencing platform or sequencing depth. While the overall lower recall and precision values were to be expected due to inaccuracies of the SV call set, we found the large rise in recall for complex substitutions questionable. We therefore examined several alignments of SVs that were typed as complex substitutions. We found that most of these complex substitutions are actually insertions or deletions, which would explain the high recall values. Most of the complex substitutions in nstd152 were determined by merging of experiments (optical mapping, sequence alignment and de novo assembly) and not associated to just one method. It is possible that conflicting findings between methods were thought to be caused by complex substitutions as they consist of both a deletion and an insertion. We added some concrete examples with screenshots of alignments and BLAST results of individual reads in the Supplementary Data as evidence of these findings.

### Improved SV calling with combiSV

This benchmark revealed the strengths and weaknesses of each SV calling tool for long read sequencing. With this performance data we were able to develop a tool (combiSV) that can combine the outputs of pbsv, Sniffles, NanoVar, NanoSV and SVIM into a superior SV call set, with Sniffles and pbsv as mandatory input. The VCF outputs of each tool serve as input and the minimal of supported reads for the variance allele has to be given. The complete wall time is under 1 minute and less than 1 GB of virtual memory is required. By combining the strengths of each of the 5 SV callers, we were able to eliminate distinct weaknesses and improve overall performance (Table 2). The most significant improvements were the ratio of total matches versus false positives and the accurate definement of the SV parameters. The added value of combiSV can also be seen by the sequence depth analysis (Figure 2), where combiSV has consistently the best overall performance and does not show any significant drops in recall or precision for any of the sequencing depths. The improved performance of combiSV is less pronounced by the precision and recall values of the individual SV types, which can be explained by the fact that the performance gain was mostly limited for deletions and insertions. Most importantly, combiSV also showed significant improvement for the real GIAB dataset, as it combines the highest recall from NanoSV, the highest precision from Sniffles and the accuracy from pbsv. This high recall is also achieved without NanoSV, as combiSV(3) only combines pbsv, sniffles and SVIM. The combination of all 5 callers reduced the recall and precision slightly, which is probably caused by the high number of false positives of NanoSV and NanoVar. Therefore it is not necessary to include the output of all 5 SV callers to run combiSV, although it is advised to add one additional caller besides pbsv and Sniffles.

## DISCUSSION

We developed a realistic simulated model to benchmark existing structural variation detection tools for long read sequencing. This was accomplished with Sim-it, a newly developed tool for the simulation of structural variation and long sequencing reads. Although there are several tools available that can simulate structural variation or long sequencing reads, a benchmark study to assess the accuracy of these simulators was needed. Besides Sim-it, the combination of Varsim and LongISLND (despite the aberration for the length of deletions) could also have been used for this benchmark study. We simulated in total 5 PacBio and 7 Nanopore whole genome sequencing datasets of GRCh38 with coverages ranging between 10x and 50x. With these simulations, we assessed the performance of 6 SV callers and the influence of increasing sequencing depths and read lengths.

For the majority of the datasets, Sniffles, pbsv and SVIM produced the best overall performance with a good balance between recall and precision. Sniffles has the lowest number of false positives for all datasets, yet performs significantly less for PacBio HiFi datasets with a coverage below 30x. pbsv defines the SVs the most accurate across all datasets and since it is designed for PacBio, it performs the best on this type data. NanoSV and NanoVar have high recall numbers, however at the cost of a disproportional high false positive rate (to a lesser extent for PacBio HiFi data). These findings were supported by our benchmark on the high fidelity SV call set of GIAB.

It is often assumed that higher sequencing depths and longer read lengths will improve assembly and variance calling outcomes. Yet in our benchmark, increasing sequencing depths does not guarantee improved structural variation calling. Although there was still a modest rise in recall numbers for sequencing depths above 30x, we did observe a disproportional rise in false positives above 30x. This rise in false positives was not observed for increasing sequencing lengths, while we observed an increase in recall for longer read lengths across all methods.

Finally, we looked at precision and recall rates for each type of SV. Each tool showed the best performance for deletions and insertions, which are the majority of SVs in a human genome. More problematic SVs are inversions and complex substitutions, wherefore recall rates are respectively between 45-65% and 1-20%. As complex substitutions are not defined by any of the tools, it seems likely that these algorithms are not designed to detect this type of SV. New SV callers or updates of existing ones could make significant improvements in this direction. Although the SV study we used as blueprint (3) detected around 3000 complex substitutions per individual, we discovered that most of these complex substitutions were insertions or deletions. The actual prevalence of this type of structural variation is therefore possibly not accurate and requires further studies in order to map the complete structural variation profile in the human genome.

This extensive benchmark unveiled the strengths and weaknesses of each SV detection algorithm and provided the blueprint for the integration of multiple algorithms in a new SV detection pipeline, namely combiSV. This Perl script can combine the VCF outputs from Sniffles, pbsv, NanoVar, NanoSV and SVIM into a superior call set that has the low false positive rate of Sniffles, the accuracy of pbsv and a high recall as SVIM. The added value of combiSV on simulated data was supported by the real dataset of GIAB, where the gains were even more outspoken.

This study shows that a simulated model can be beneficial to gain a better understanding in the performance of structural variation detection tools. It is crucial that the simulations are as accurate as possible. Currently, Sim-it does not simulate small indels and SNPs, although they can have an effect on the detection of small SVs and will therefore be included in the next update. The sequencing depth of real sequencing datasets show much more fluctuations than a simulated one, we therefore propose to include a profile of the sequencing depth in a real dataset that can be reproduced for the simulation.

## METHODS

### Sim-it

We developed a new structural variation and long read sequencing simulator, called Sim-it. The structural variation module outputs fasta files of each haplotype, plus an additional one that combines all SVs in one sequence. A set list of SVs can be combined with additional random generated SVs as input. The long read sequencing module outputs sequencing reads based on a given error profile and 4 metrics (coverage, median length, length range and accuracy). We provide error profiles for Nanopore, PacBio RS II, Sequel II and Sequel HiFi reads. Additional error profiles can be generated with a custom script. Both simulation modules (SV and long reads) can be used separately or simultaneously, starting from a sequence file as input. We also provide plots with the length distributions for the simulated sequencing reads and structural variations (insertions, deletions and inversions). Sim-it was written in Perl and does not require any further dependencies. Sim-it is open source and can be downloaded at https://github.com/ndierckx/Sim-it, where a more complete manual can be found.

### Benchmark of structural variation simulators

We compared Sim-it (v1.0) with RSVSim (v1.24.0) (13), SVEngine (v1.0.0) (14), SCNVSim (v1.3.1) (10) and VarSim (v0.8.4) (11) for computing resource consumption and available features. Runtime performance was measured using the Unix time command and Snakemake (v5.7.0) (15) benchmark function on the custom VCF of 24,600 SVs. We did not evaluate SCNVSim performance because it does not accept a custom VCF file. All scripts were executed on a Xeon E7-4820 with 512GB of memory.

### Benchmark of the long read simulators

We compared Sim-it (v1.0) with the long read simulators PBSIM (v1.0.4) (16), Badread (v0.1.5) (17), PaSS (18), LongISLND (v0.9.5) (19), DeepSimulator (v1.5) (20), Simlord (v1.0.3) (21) and NanoSim (v2.6.0) (22) for computing resource consumption and error frequency within context-specific patterns for mismatches and indels using real data of Nanopore and PacBio sequencing. Runtime performance was measured using the Unix time command and Snakemake (v5.7.0) benchmark function on the 15x sequencing coverage simulation with chromosome 1 of GRCh38. Context-specific error patterns were analyzed by a custom perl script with alignment 30x simulated read to 60 Kbp sequence. All scripts were executed on a Xeon E7-4820 with 512GB of memory. More details on the error profiles used for each simulation can be found in the Supplementary Data.

### Train customized error profiles for Sim-it

The E. coli K12 substrain MG1655 dataset of PacBio Sequel II and PacBio RS II was downloaded from the github website of Pacific Biosciences. Using the above two datasets we trained the error profile of PacBio Sequel II and PacBio RS II. We also downloaded the GIAB HG002 dataset of PacBio Sequel II HiFi reads powered by CCS. To reduce the computational time, we trained the error profile of PacBio Sequel II HiFi reads based on chromosome 1 of GRCh38. The Nanopore error profile is based on sequencing reads of a human sample on PromethION 9.4.1 flow cells.

### SV detection on simulated reads

We used the simulated data from Sim-it to validate 6 structural variant callers, namely Sniffles (v1.0.11) (1), SVIM (v1.3.1) (23), NanoSV (v1.2.4) (24), Picky (v0.2.a) (25), NanoVar (v1.3.8) (26) and pbsv (v2.3.0). A list of 24,600 SVs, derived from sample NA19240 of dbVAR nstd152 (3), was used to simulate Nanopore, PacBio CLR reads and PacBio HiFi reads for GRCh38 at a sequencing depth of 20x. We also simulated 20x normal read using GRCh38 with not structural variants at all. Besides for pbsv, we aligned the simulated reads to GRCh38 using Minimap2 (v2.17-r941) (27). The alignment for pbsv was performed using pbmm2 (v1.3.0) with default parameters. The exact parameters that were used for the alignments and SV callers can be found in the Supplementary Data.

Furthermore, we simulated additional Nanopore and PacBio HiFi reads for GRCh38 at sequencing depths of 10x, 30x and 50x to study the influence of increasing sequencing depths for SV calling. Each of the Nanopore simulations had a median read length of 25,000 bp, we also included two additional simulations of 15,000 bp and 40,000 bp with a sequencing depth of 20x. PacBio long reads have a median length of 25,000 bp and the PacBio HiFi reads a median length of 15,000 bp. An additional filtering step was added for each VCF output; we only retained variances that obtained a PASS for the FILTER value, that have a length of 50 bp or more and wherefore at least 3 (for sequencing depths 10x and 20x) or 5 (for sequencing depths 30x and 50x) reads support the variance. This additional filtering step significantly improved the output for each tool compared to the raw VCF output.

Benchmark metrics were calculated by comparing the VCF output of each SV caller against the simulated reference set of 24,600 SVs. For each detected SV, we looked for possible matches in the reference set within a 500 bp range of the detected position. When the length of the SV was determined, we tolerated an error margin of 30%. If these two conditions were met, a detected SV was matched to the SV of the reference set, independent from the type or haplotype that was called. As there are multiple metrics that define the performance of an SV detection algorithm, we adopted an overall score that that combines each of the metrics. For each detected SV, a maximal score of 1 was possible; 0.4 for the correct position, 0.2 for the correct length, 0.2 for the correct type of SV and 0.2 for the correct haplotype. The scores for length and position proportionally decreased with difference compared to the reference set. Finally, the number of false positives were subtracted from the total score and eventually expressed as a percentage of the maximum possible score (Table 2).

### SV detection on real datasets

The Genome in a Bottle (GIAB) Consortium recently developed a high-quality SV call set for the son (HG002/NA24385) of a broadly consented and available Ashkenazi Jewish trio from the Personal Genome Project. We performed a benchmark on the latest most conserved BED file (HG002_SVs_Tier1_v0.6.2.bed) for this sample, which contains 5,260 insertions and 4,138 deletions.

The public available ultralong Nanopore reads (GM24385) with an average sequencing depth of 45x were used for this benchmark. Furthermore, we compared SV detection metrics of a public available PacBio dataset of NA19240 (3) with an average sequencing depth of 37x against the results of our simulated datasets.

### combiSV

With the results of the SV detection benchmark, we developed a script to combine the results of pbsv, Sniffles, NanoVar, NanoSV and SVIM. The output VCF files of each of the 5 tools serve as input, from which the files of pbsv and Sniffles are obligatory to run combiSV. The minimal coverage of the alternative allele is set to 3 as default value, but can be adjusted for datasets with high sequencing depths. The script was written in Perl and does not require any further dependencies. combiSV is open source and can be downloaded at https://github.com/ndierckx/combiSV.

## ACKNOWLEDGEMENTS

This project was supported by FWO TBM project T003819N to J.R.V. We thank the Center for Precision Medicine at Sun Yat-sen University for providing the high performance computers. This project was supported by National Key R&D Program of China (2019YFA0904401 to Z.X.)

## AUTHOR CONTRIBUTIONS.

ND conceived, designed and scripted Sim-it and combiSV. ND analyzed the data for the SV caller benchmark. TL ran the simulations and existing SV calling tools for the benchmark. TL and ND designed and executed the SV and long read simulator benchmark. ND wrote the manuscript. JRV and ZX provided guidance and reviewed the manuscript.

