## Supplementary Materials for "A benchmark of structural variation detection by long reads through a realistic simulated model"

### Parameters for the structural variation callers

1. Sniffles (v1.0.11)

We aligned the simulated reads to GRCh38 using Minimap2 (v2.17-r941). We changed the parameters of Minimap2 according to the type of simulated reads, 'minimap2 -ax map-ont' corresponds to Nanopore simulated reads and 'minimap2 -ax map-pb' corresponds to PacBio simulated reads.
The parameters of Sniffles is 'sniffles -n -1 -s 3 --genotype'.

2. SVIM (v1.3.1)

The same alignment as for Sniffles. We used SVIM to call structural variants with default parameters.

3. NanoSV (v1.2.4)

The same alignment as for Sniffles. We used NanoSV to call structural variants without 'depth_support' mode in its config file. Other parameters are default.

4. Picky (v0.2.a)

The same alignment as for Sniffles. We used the pipeline provided by the github page of Picky to call SVs (https://github.com/TheJacksonLaboratory/Picky/wiki/Using-an-Alternative-Aligner).

5. NanoVar (v1.3.8)

We changed the parameters of NanoVar according to the type of simulated reads, 'nanovar -l 50 -x ont' corresponds to Nanopore simulated reads, 'nanovar -l 50 -x pacbio-clr' corresponds to PacBio simulated reads and 'nanovar -l 50 -x pacbio-ccs' corresponds to PacBio HiFi simulated reads.

6. pbsv (v2.3.0)

We aligned simulated reads to GRCh38 using pbmm2 (v1.3.0) with default parameters. The parameters of pbsv is 'pbsv call -m 50'.

### Long read simulators benchmark

We compared the features of 8 different long read simulators. We tested the wall time and memory consumption for the simulation of 15x coverage Nanopore or PacBio (when no Nanopore available) long reads for the human chromosome 1 (GRCh38).

**Table 1 |** Comparison of the features of each long read simulator and a system requirements benchmark.

| | Sim-it | PBSIM | Badread | PaSS | LongISLND | DeepSimulator | Simlord | NanoSim |
|---|---|---|---|---|---|---|---|---|
| Error profiles | ONT, PB (RS2, Sequel2, Sequel HiFi) | PB (CCS, RS) | ONT, PB (RS2) | PB (Sequel, RS2) | ONT, PB (RS2) | ONT | PB (CCS, RS2) | ONT |
| Train model | ✔ | | ✔ | ✔ | ✔ | ✔ | | ✔ |
| Accuracy adjustment | ✔ | ✔ | ✔ | | ✔ | | ✔ | |
| Read length adjustment | ✔ | ✔ | ✔ | | ✔ | ✔ | ✔ | ✔ |
| Transcriptome reads | | | | | | | | ✔ |
| Separate haplotypes | ✔ | | | | | | | |
| Quality scores | | ✔ | ✔ | ✔ | ✔ | ✔ | ✔ | |
| Wall time | 64 min | 5 min | 938 min | 12 min | 31 min | 1634 min | 81 min | 327 min |
| Virtual Memory | 1,2 GB | 0,25 GB | 12,1 GB | 2,5 GB | 2,5 GB | 14,8 GB | 0,37 GB | 0,43 GB |

### Detail for error profile of long read simulators

The error profile of PBSIM (v1.0.4) is located in the installation folder and the relative path is 'PBSIM/data/model_qc_clr'. The error profile of Badread (v0.1.5) was set '--error_model nanopore' and '--error_model pacbio' for the respective simulations. The error profile of PaSS is located in the installation folder ('PaSS/E.coli/ecoli.config'). We used '-m pacbio_sequel -c PaSS/E.coli/ecoli.config' for the simulation of PaSS. LongISLND (v0.9.5) does not provide an error profile so we trained error profiles using the same datasets as with Sim-it (v1.0). DeepSimulator (v1.5) provides a Nanopore error profile in the installation folder. Simlord (v1.0.3) does not provide error profiles so we changed the parameters 'Probability for insertions, deletions, substitutions' according to the observed values of real PacBio datasets. The command is 'simlord -ps 0.0312 -pd 0.0309 -pi 0.0433'. We downloaded the NanoSim (v2.6.0) error profile named 'human_NA12878_DNA_FAB49712_albacore' from it's website.

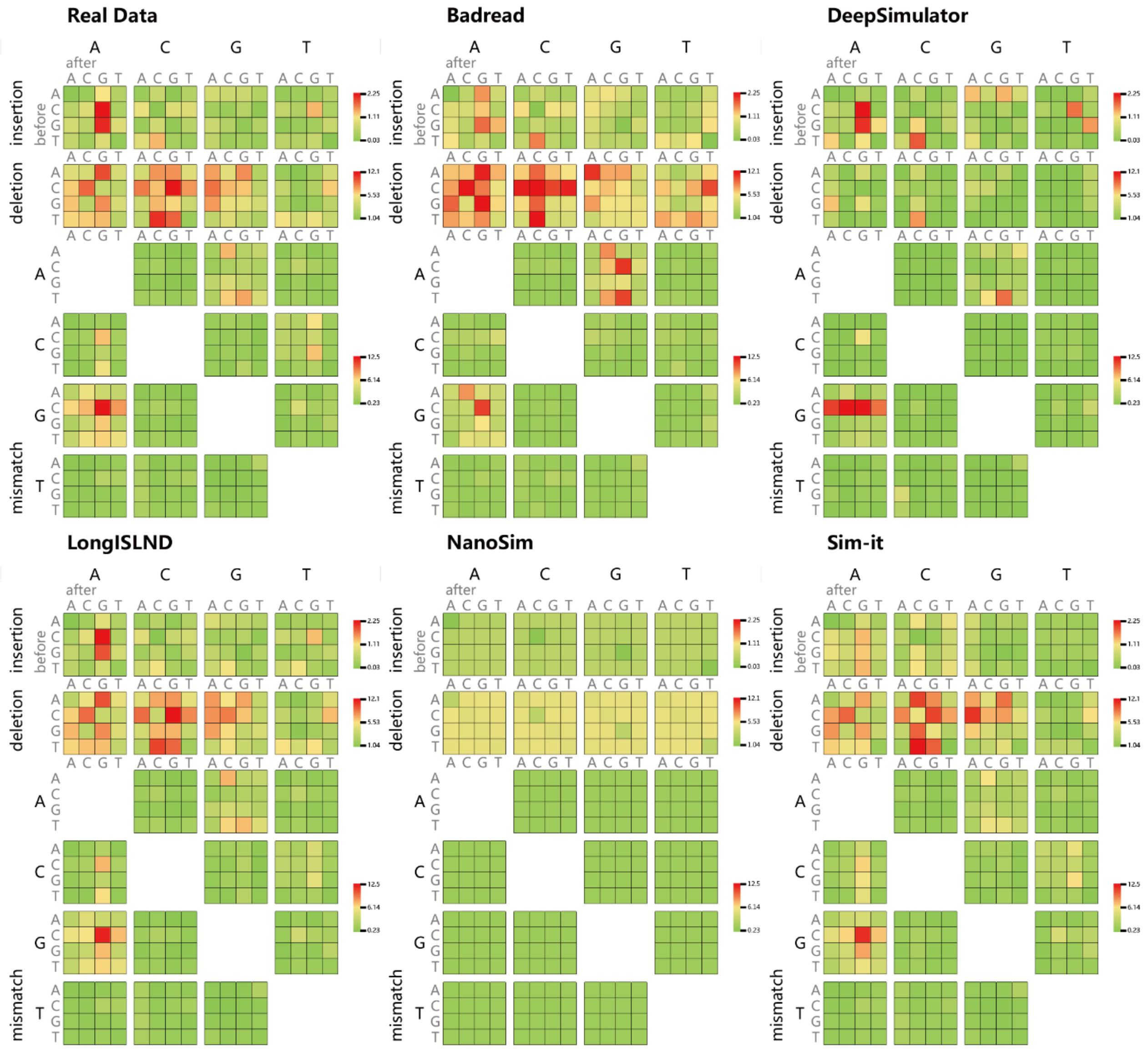


**Figure 1 |** Error profiles of simulated Nanopore reads from 5 different long read simulators.

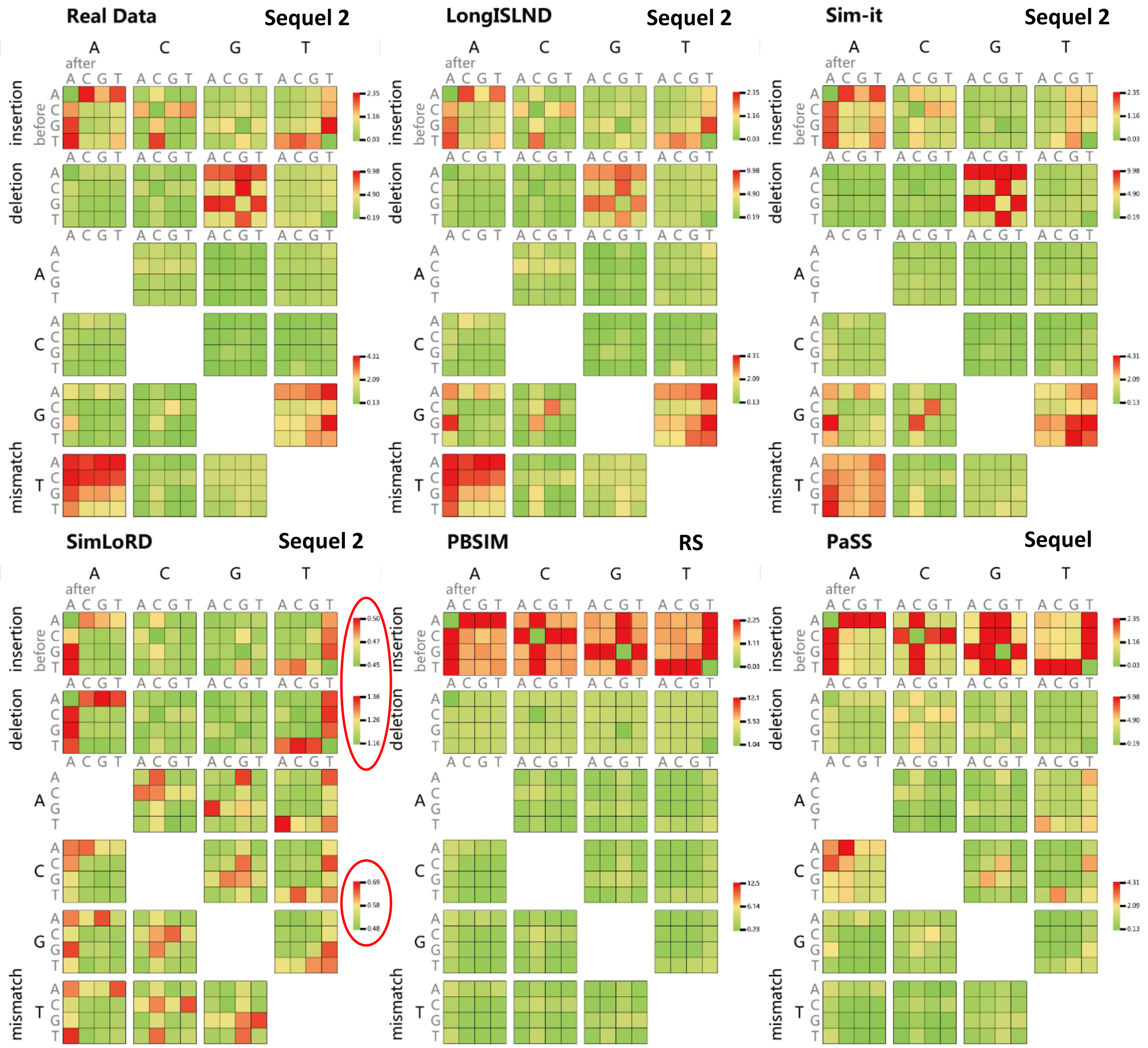


**Figure 2 |** Error profiles of simulated PacBio reads from 5 different long read simulators. The error rate for Simlord is much lower compared to the real data, we therefore adjusted the ratios to visualize the error profile.

### Complex substitutions in NA19240

The recall of complex substitutions (CSUB) are significantly higher for the real PacBio dataset (60%) of NA19240 than for our simulated datasets (1-20%). Because we expected a drop in recall we examined the alignment of 27 CSUBs manually with IGV. We selected only homozygous CSUBs to simplify the interpretation of the alignments. CSUBs were not selected on any other criteria, we selected 12 random homozygous CSUBs from chromosome 1 and 15 random homozygous CSUBs of chromosome 2. For the simulated CSUBs, the length of the deleted sequences were always the same as the length of the inserted sequences. This is not necessarily the case for the real CSUBs, which could partially explain the discrepancy between the recall values and different alignment patterns. Nevertheless, several of the CSUBs we examined were in fact deletions or insertions that were incorrectly categorized as a CSUB. From only examining the alignments, we could only confirm one CSUB out of the 27 potential CSUBs as a true CSUB. For 6 presumed CSUBs, we aligned and inspected several individual PacBio reads separately. When the alignment does not show any SV at the given position, it also possible that called position was inaccurate. For each of the screenshots of IGV, the top alignment is the simulation and the bottom the true dataset of NA19240.

Figure 3: We did not observe a SV at this position by inspecting the alignments.

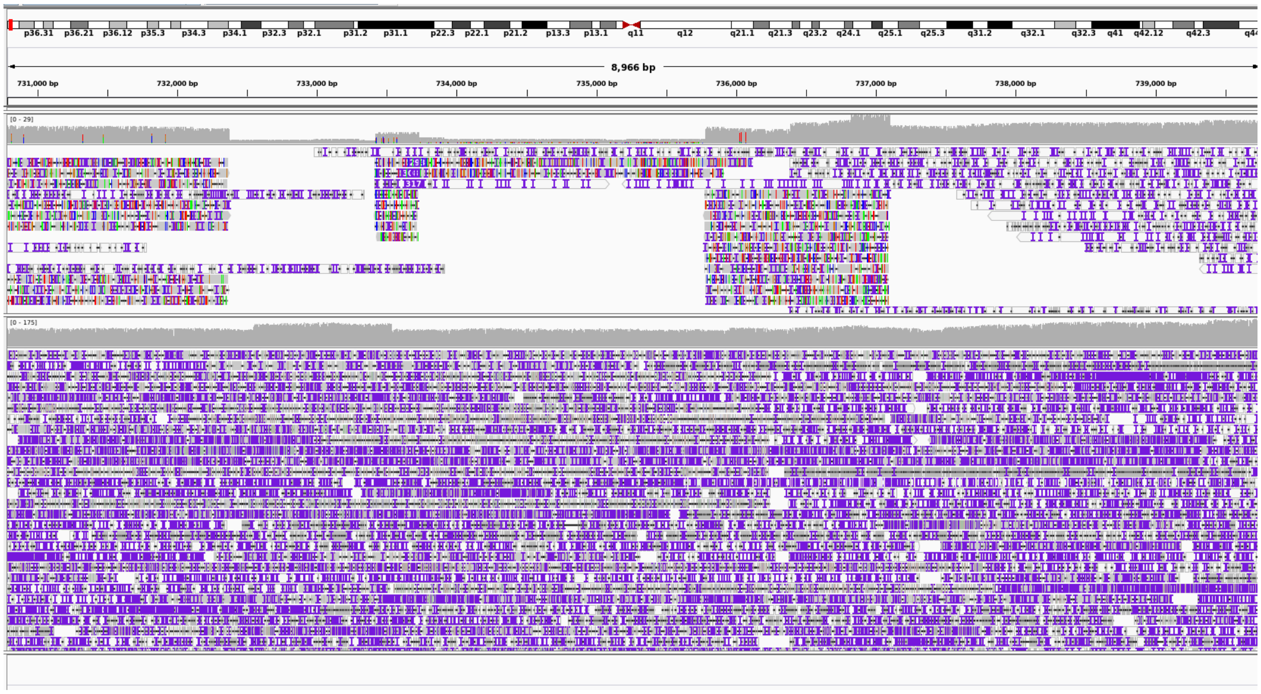


**Figure 3 |** CSUB chromosome 1, position 732377, length 3402 bp

Figure 4: This is the only alignment that visually resembles a theoretical CSUB. SV callset ‘nstd152’ called 3 CSUBs of 61 bp within a region of 400 bp. This region is a tandem repeat region and when we individually aligned several PacBio reads, we found that there are two different haplotypes. Both haplotypes have a shorter tandem repeat region compared to the reference and for one haplotype we also observed an insertion of around 100 bp and can there be categorized as a heterozygous CSUB.

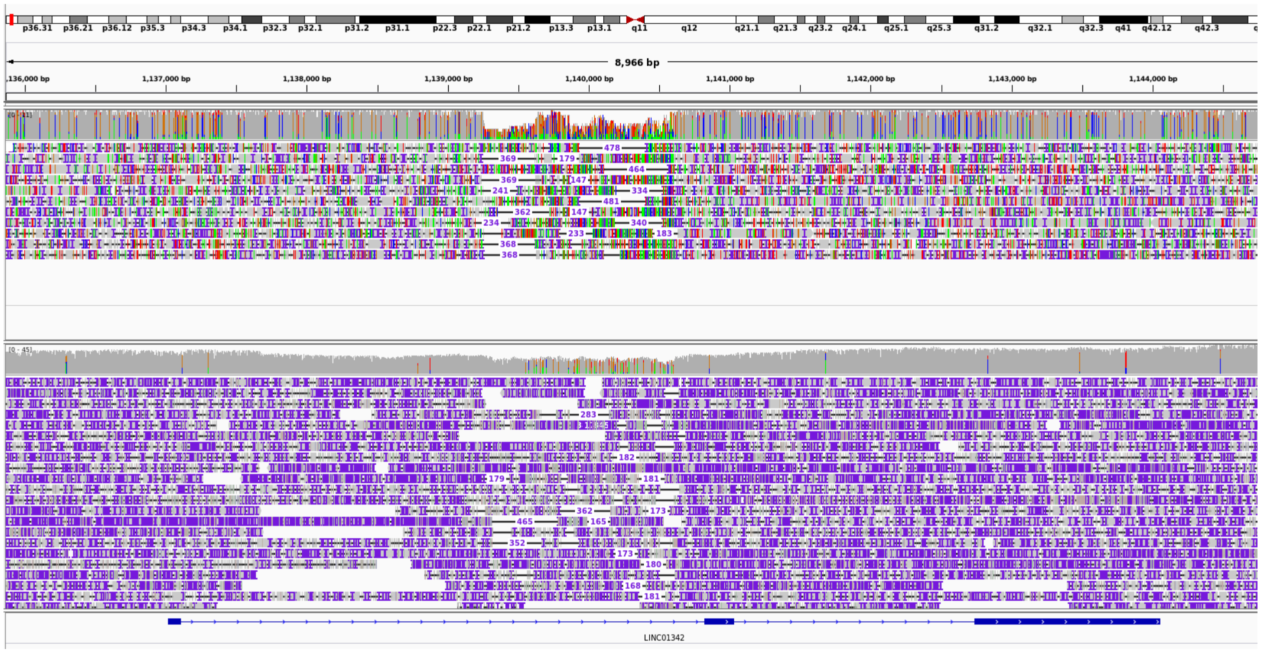


**Figure 4 |** CSUB chromosome 1, position 1140181, length 61 bp

Figure 5 & 6: This SV is a confirmed deletion, as can be seen in the overall alignment and the individual PacBio alignments.

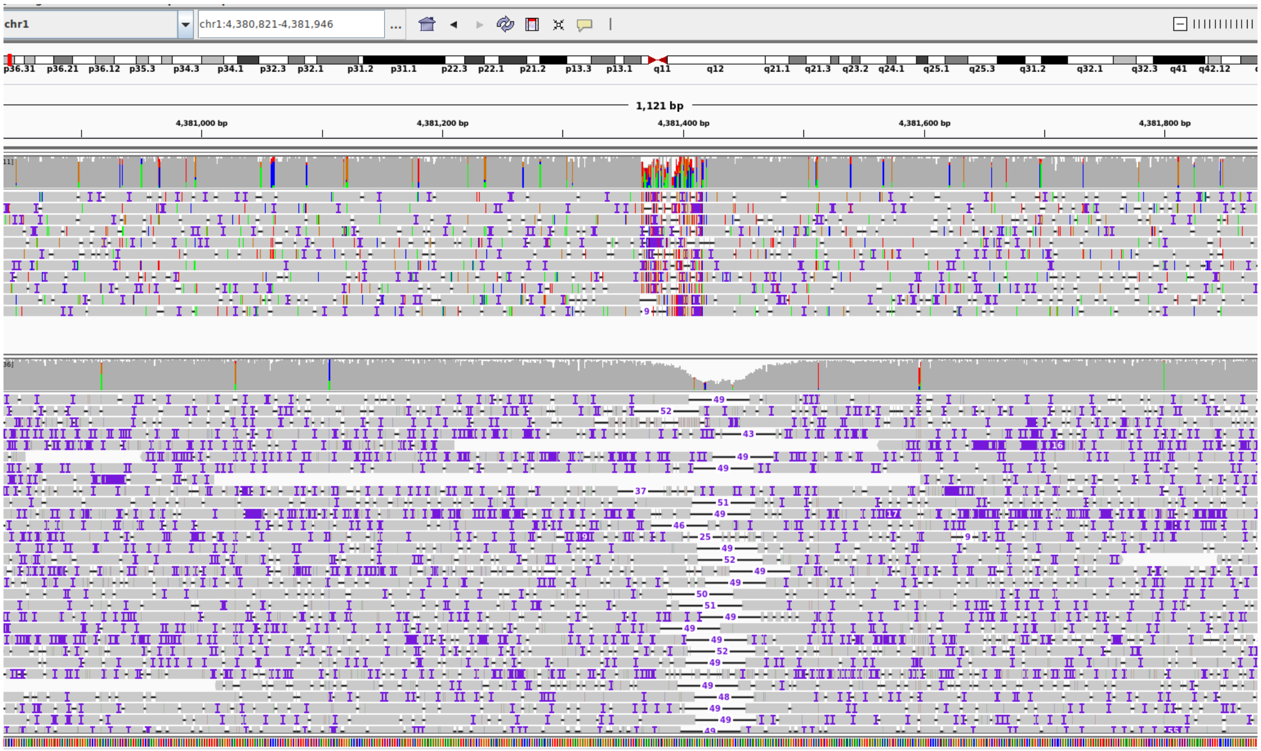


**Figure 5 |** CSUB chromosome 1, position 4381366, length 51 bp.

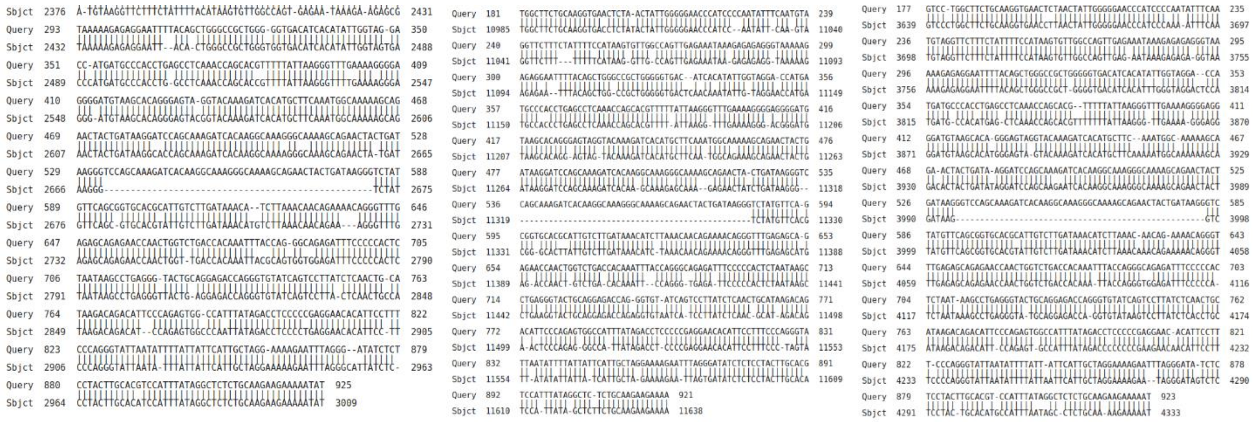

**Figure 6 |** Individual alignment of three PacBio reads for ‘CSUB chromosome 1, position 4381366, length 51 bp’.

Figure 7: Several reads show insertions around this position.

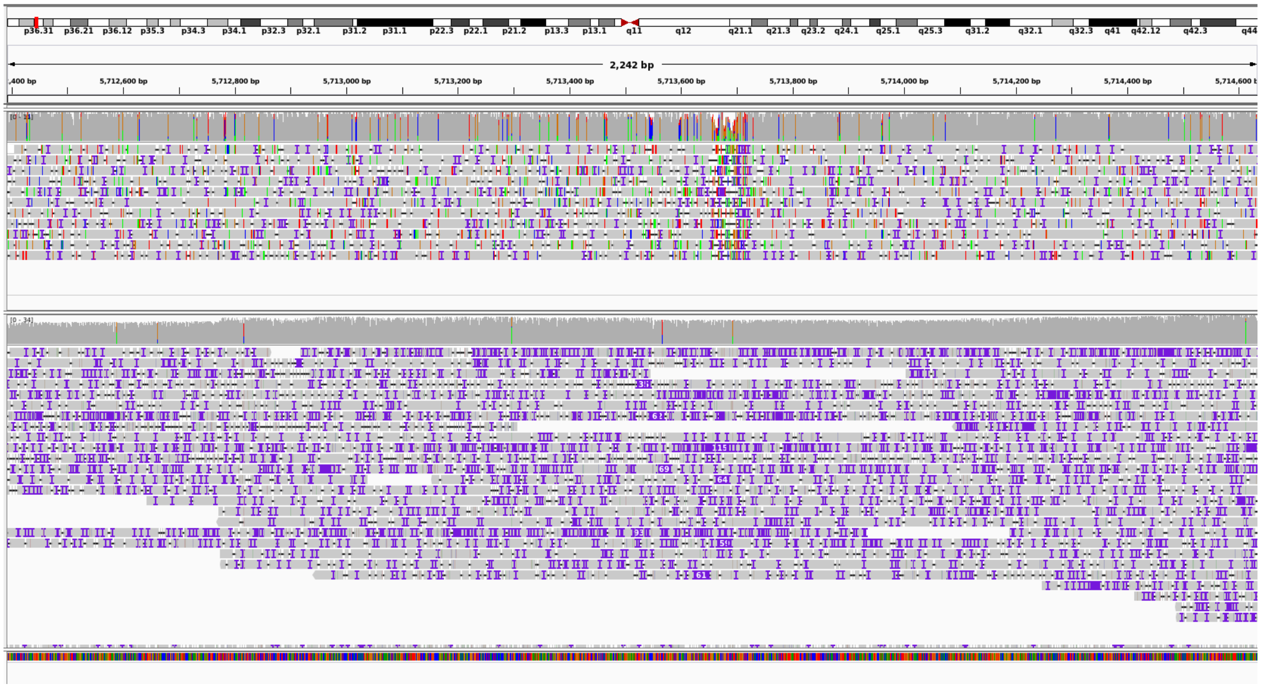


**Figure 7 |** CSUB chromosome 1, position 5713658, length 62 bp

Figure 8 & 9: We did not observe a SV at either position by inspecting the alignments.

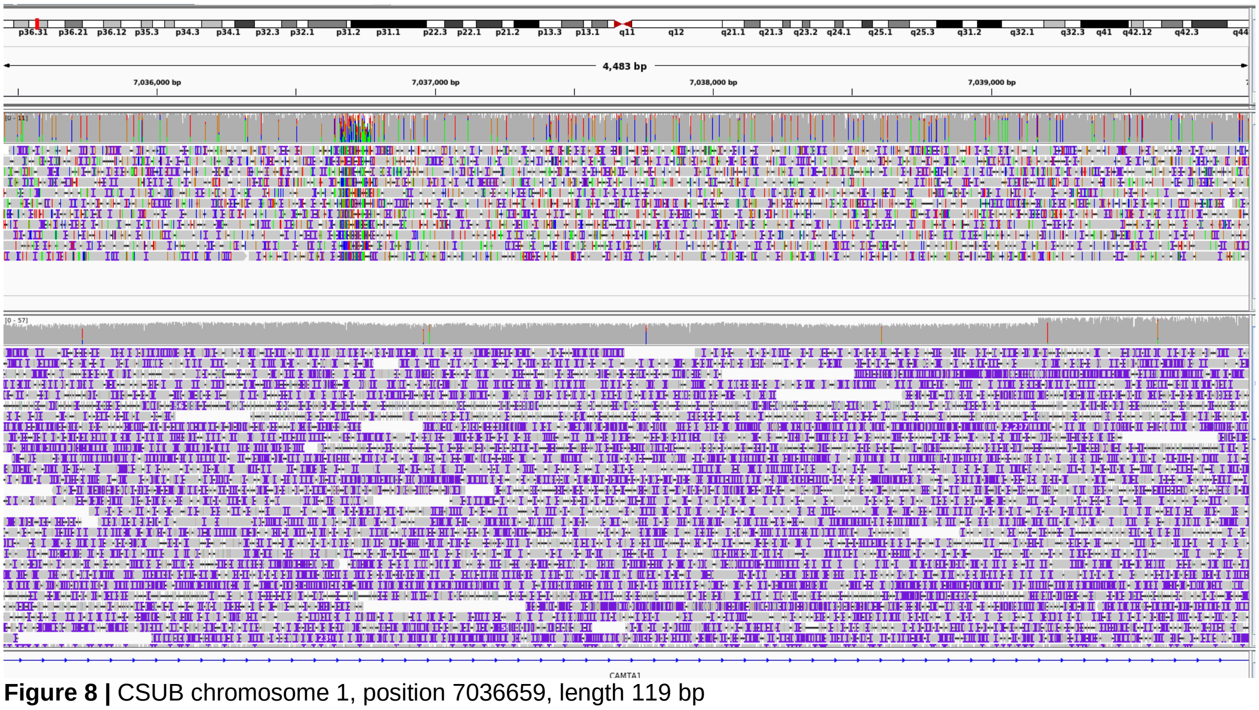


**Figure 8 |** CSUB chromosome 1, position 7036659, length 119 bp

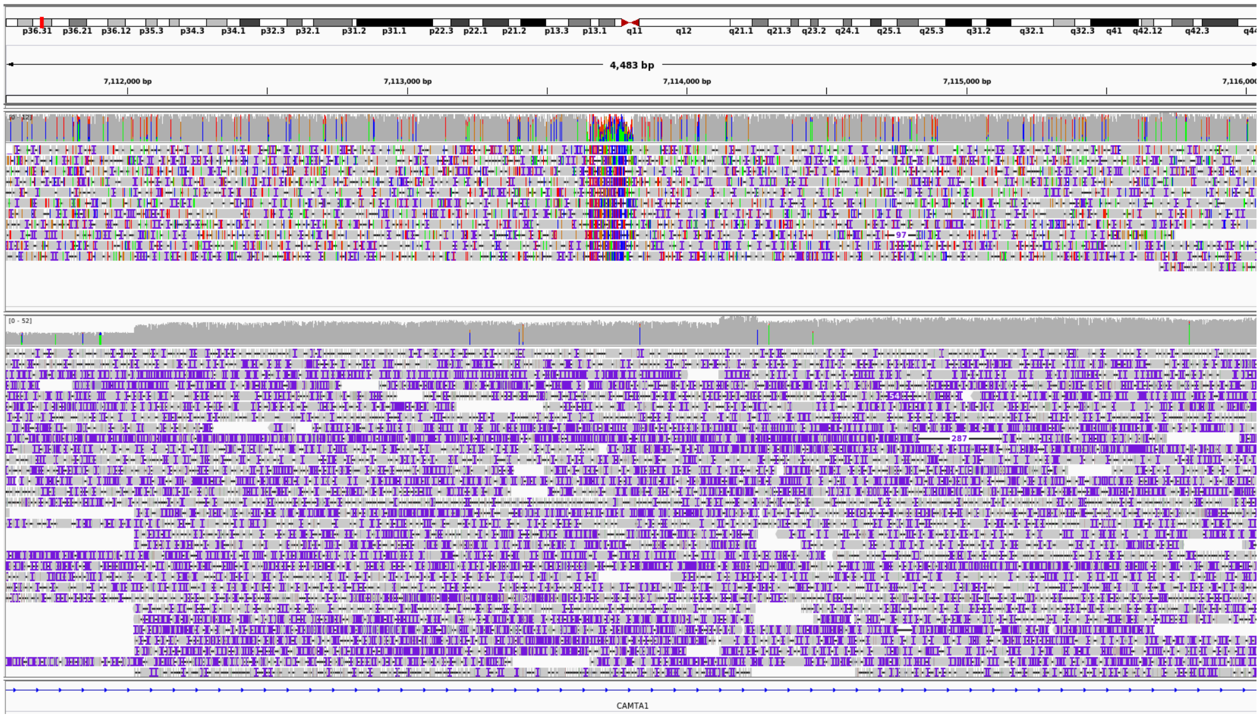


**Figure 9 |** CSUB chromosome 1, position 7113644, length 165 bp

Figure 10 & 11: This SV is a confirmed insertion, as can be seen in the overall alignment and the individual PacBio alignments.

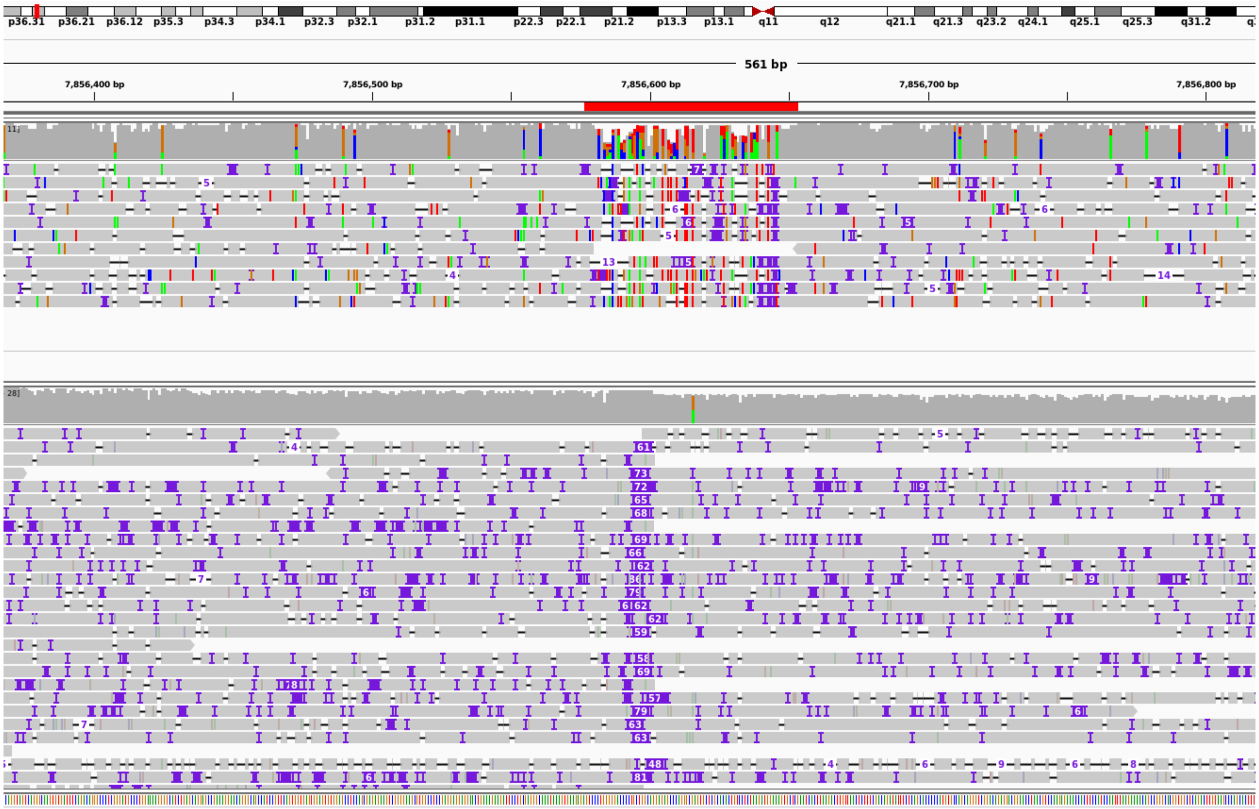


**Figure 10 |** CSUB chromosome 1, position 7856583, length 62 bp

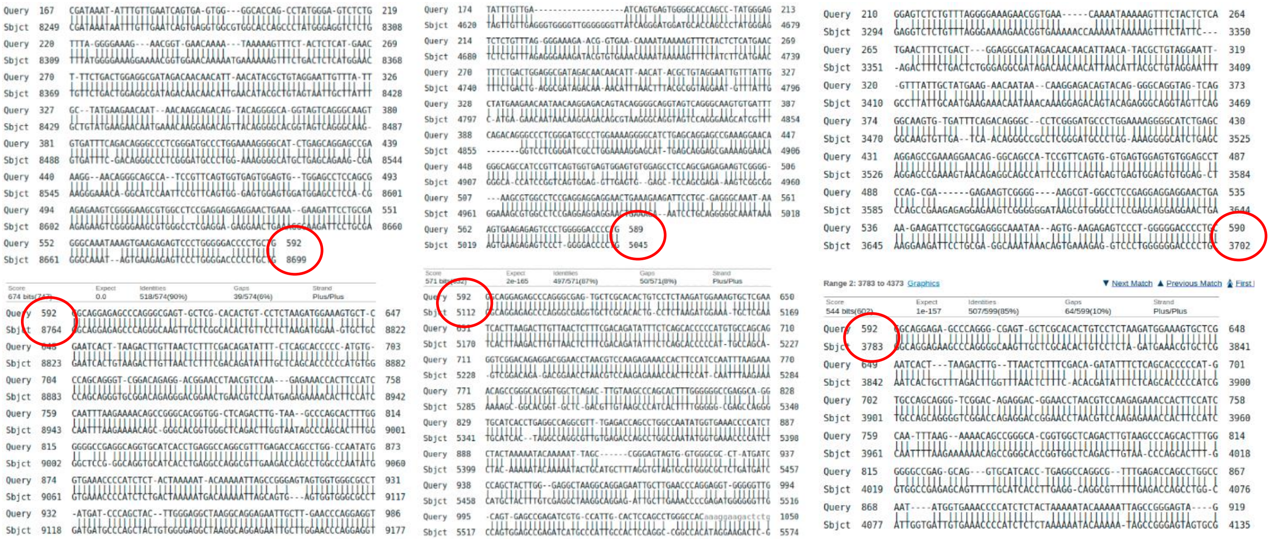

**Figure 11 |** Individual alignment of three PacBio reads for ‘CSUB chromosome 1, position 7856583, length 62 bp’.

Figure 12 & 13: This SV is a confirmed deletion, as can be seen in the overall alignment and the individual PacBio alignments. By comparing individual PacBio reads with the reference we concluded that the first duplicated sequence was deleted in the NA19240 (first yellow sequence in Figure 13).

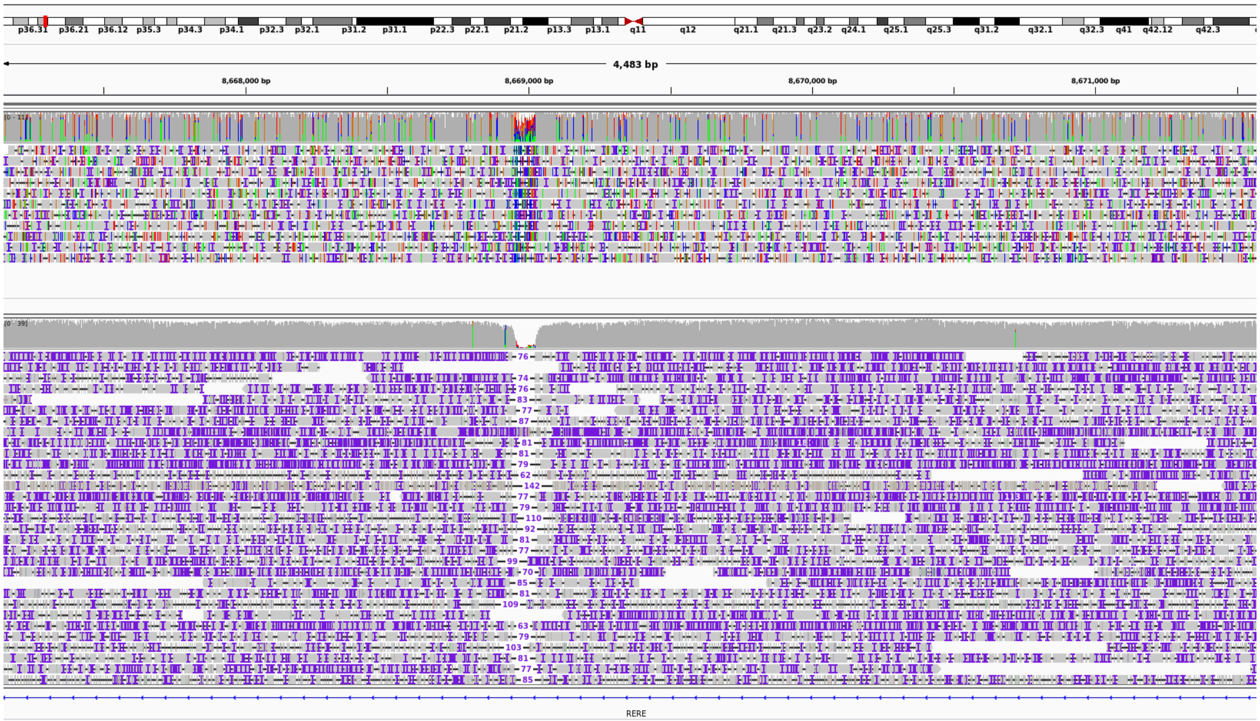


**Figure 12 |** CSUB chromosome 1, position 8668948, length 76 bp

REFERENCE
ACTAAACTCTGTGTGTGTGTGTGTGTGTGTGTGTGTGTGTGTGTGTGTGTGTGTTGTGTTTTTAAGACAGGGTCTCACTCTGTCTTCTGTGAATATTCTAAAATGCACTCAACTCTGTGTGTGTGTGTGTGTGTGTGTGTGTGTGTGTGTGTGTGTGTGTGTGTGTGTGTGTTGTGTTTTTAAGACAGGGTCTCACTCTGTCACCCA

READ1
ACTCAACTTTGTGTGTGTGTGGTGTGTGTGTGGTGTGTGTGTGGTTTGTGGTGTGTGTTGTGTGTGTGTGGGTGTGTTGTGGTTTTAAGACAGGGTCTCACTCTGTCACCCAGGCTGGGGTGCAGTGGCATGCGTGATCACGGCTCACTGTCAGCCTTGACCTACCGGGCTCAAGCATACCCCCAACTTCGCCT

READ2
ACTCAACTCTGGTTGTGTGTGTGTGTGTTGTGTGTGTGTTGTGTGTGGTGTGTGTGTTGTGTGTGTGTGTGGTTTTAAGACAGGGTCTCACTCTGTCACCAGGCTGGAAGTGCAGTGCCATGCGTGAATCACGGCTCACTGCAGCCTTGACCTATTCGGCTCAAGCAATACCCCCACTCGCCTCTCTCGTAGCTG

READ3
CTCCAACTCTGTTGTGTGTGTGTGTGTGTGTGTGTGTGTGTGTGTGTGTGTGTGTGTGTGTGTGTGTGTGTTGGTGTGTGTTGTGTTTGTGTTTTTAAGACAGTCTCACTCTGCTCACCCAGGCTGGAGTGCAGCTGCGAACATGCGCTGATCACGGCTCACTGCAGCCTTGACCTACCGGGCTCAAGCAATACCCCCCCAAC

**Figure 13 |** Individual alignment of three PacBio reads for ‘CSUB chromosome 1, position 8668948, length 76 bp’.

Figure 14 & 15: This SV is a confirmed insertion, as can be seen in the overall alignment and the individual PacBio alignments. Although it seems this insertion was heterozygous, as half of the reads do not show any inserted sequence.

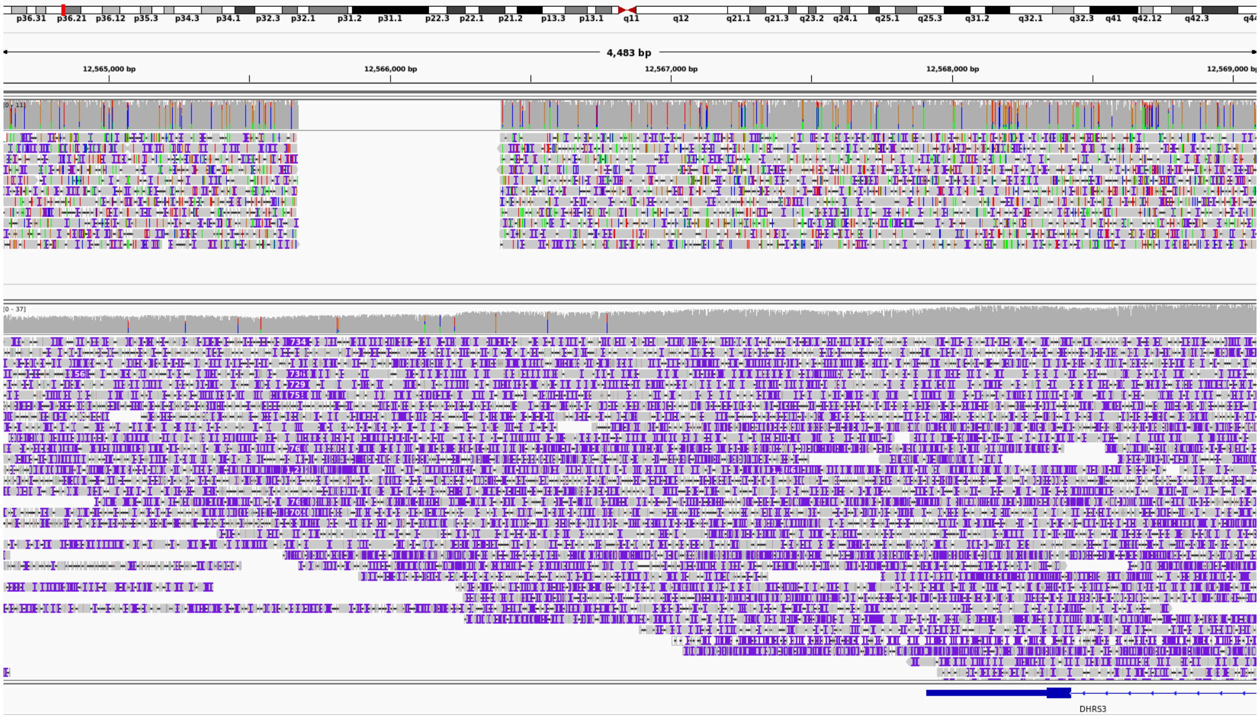


**Figure 14 |** CSUB chromosome 1, position 12565675, length 719 bp

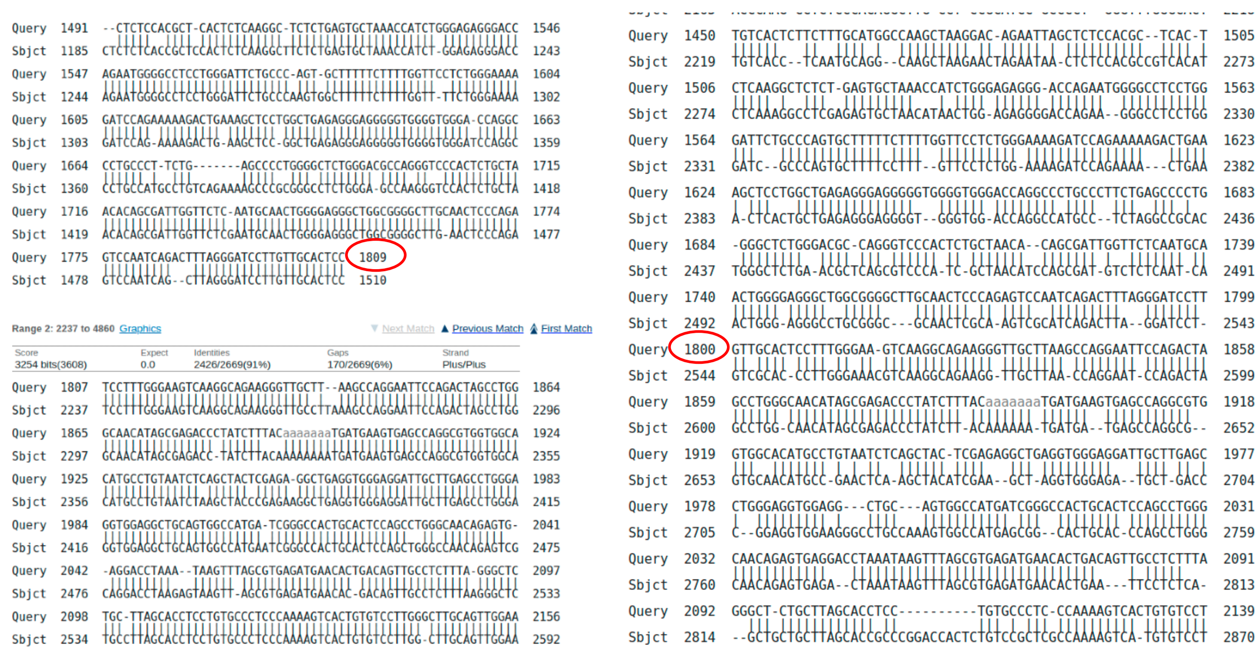


**Figure 15 |** Individual alignment of two PacBio reads for ‘CSUB chromosome 1, position 12565675, length 719 bp’.

Figure 16: Unconfirmed SV.

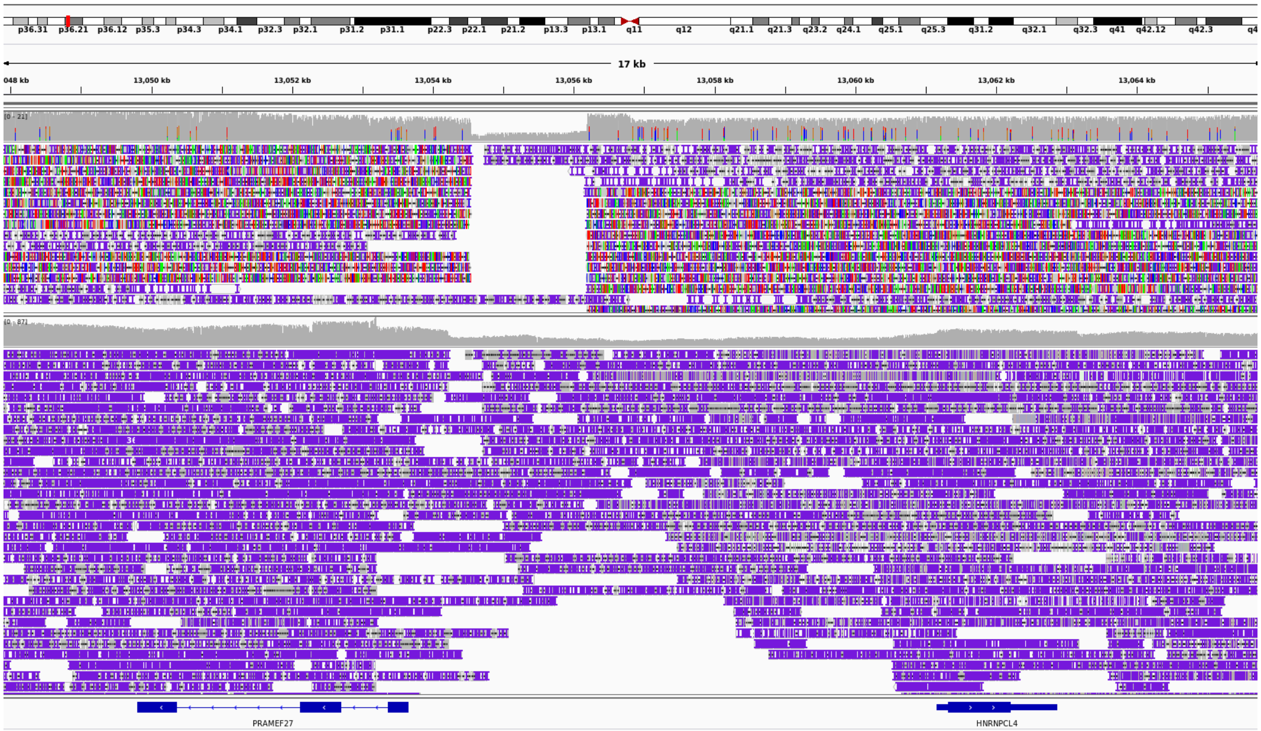


**Figure 16 |** CSUB chromosome 1, position 13054543, length 1646 bp

Figure 17: We observed an insertion, although this is a duplicated region which makes alignments less reliable.

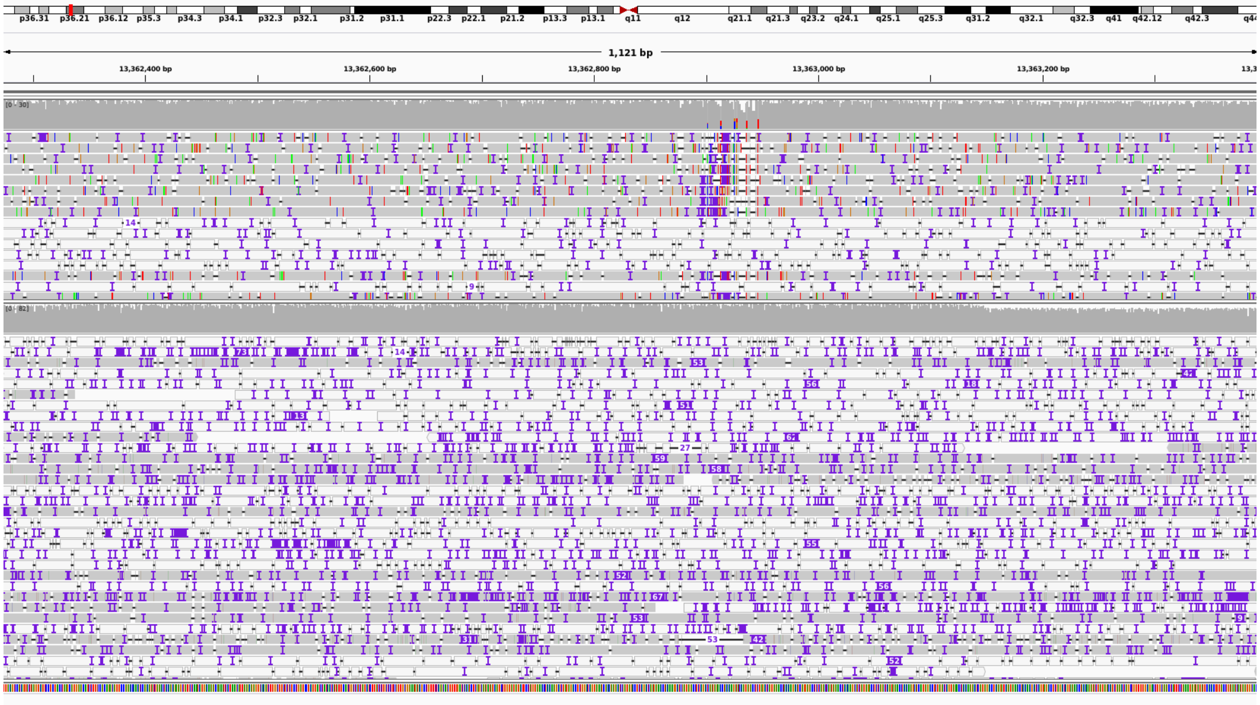


**Figure 17 |** CSUB chromosome 1, position 13362897, length 53 bp

Figure 18: We observed an heterozygous insertion.

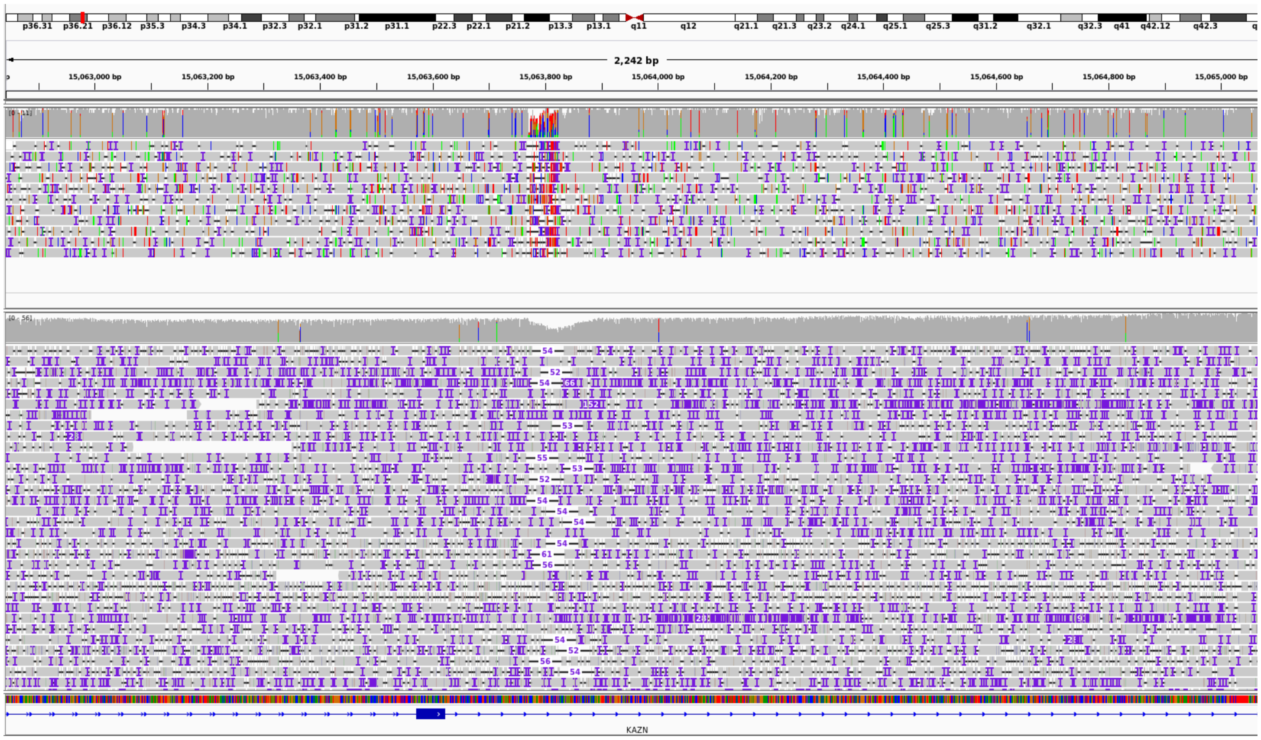

**Figure 18 |** CSUB chromosome 1, position 6935636, length 64 bp

Figure 19: We observed a deletion.

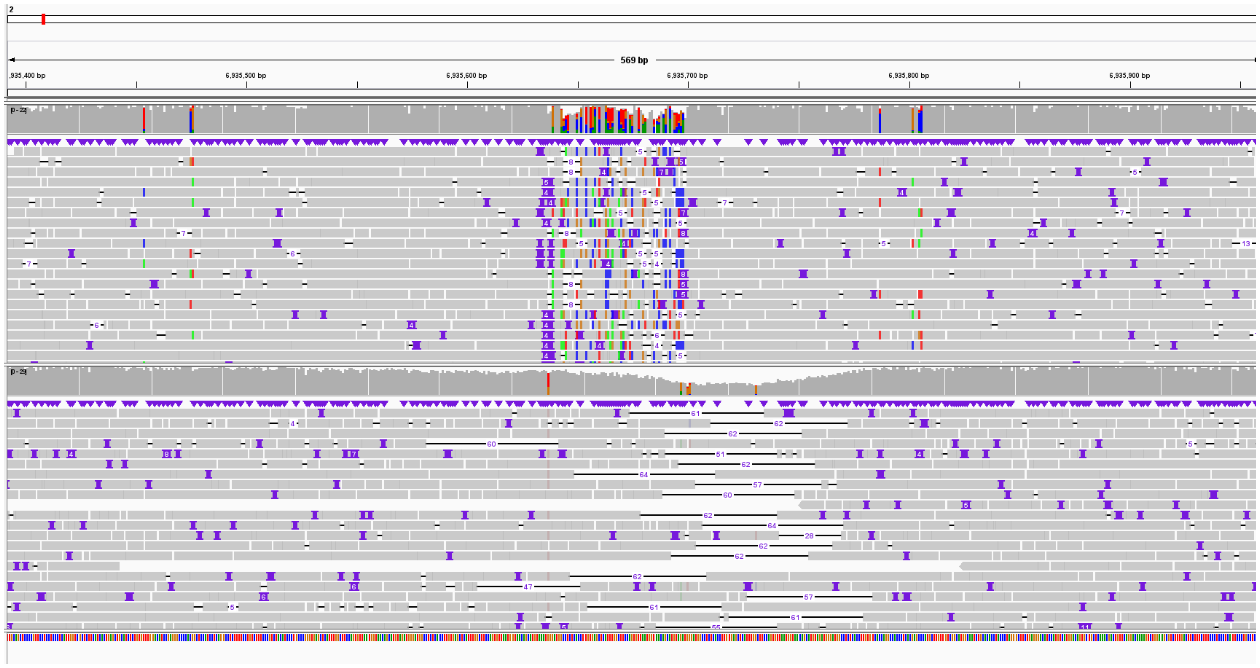

**Figure 19 |** CSUB chromosome 2, position 6935636, length 64 bp

Figure 20 - 22: We did not observe a SV at any of the positions by inspecting the alignments.

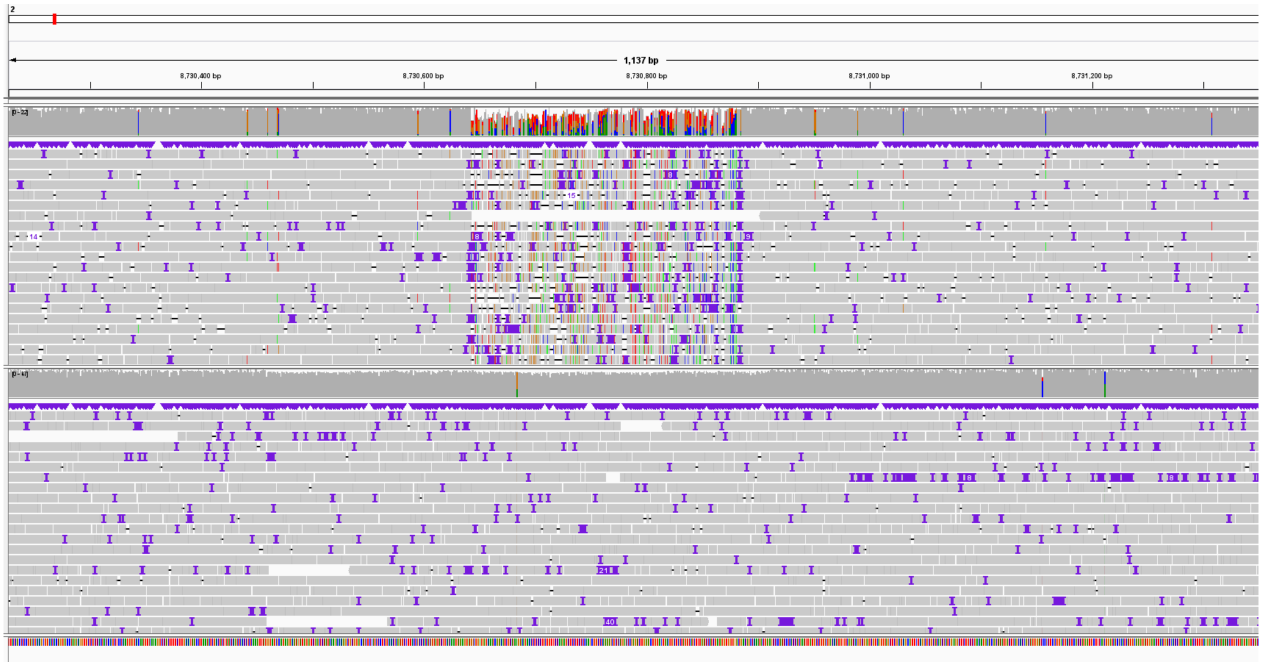


**Figure 20 |** CSUB chromosome 2, position 8730642, length 242 bp

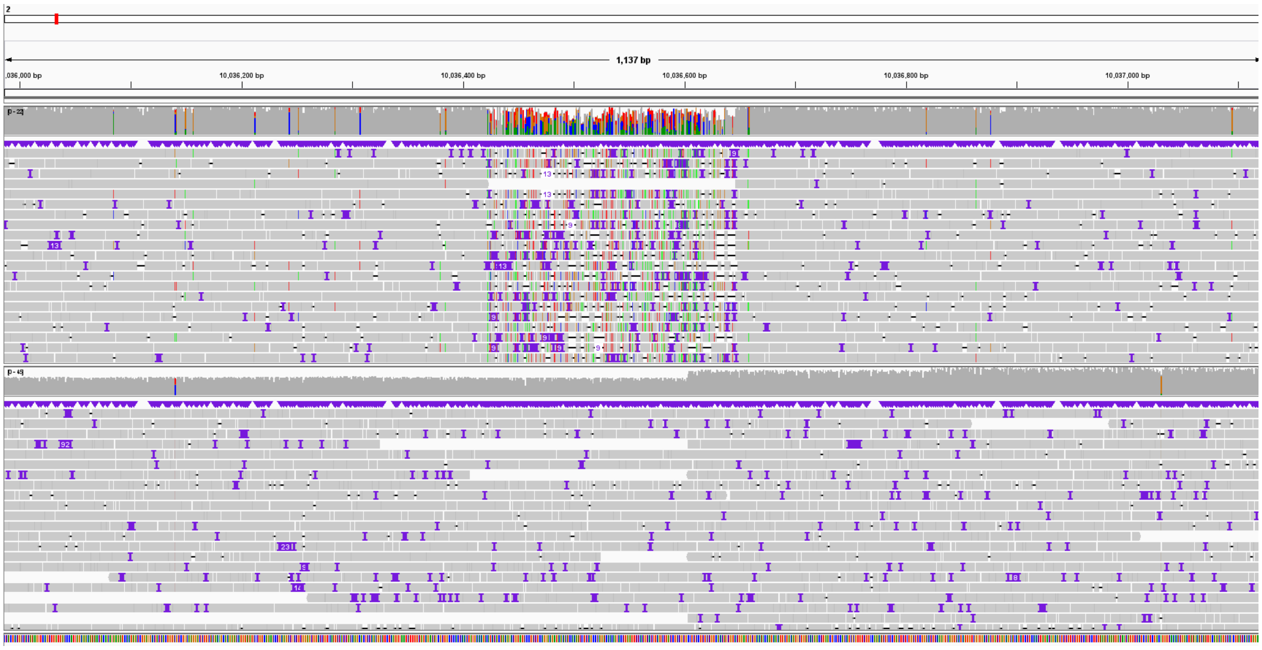


**Figure 21 |** CSUB chromosome 2, position 10036422, length 225 bp

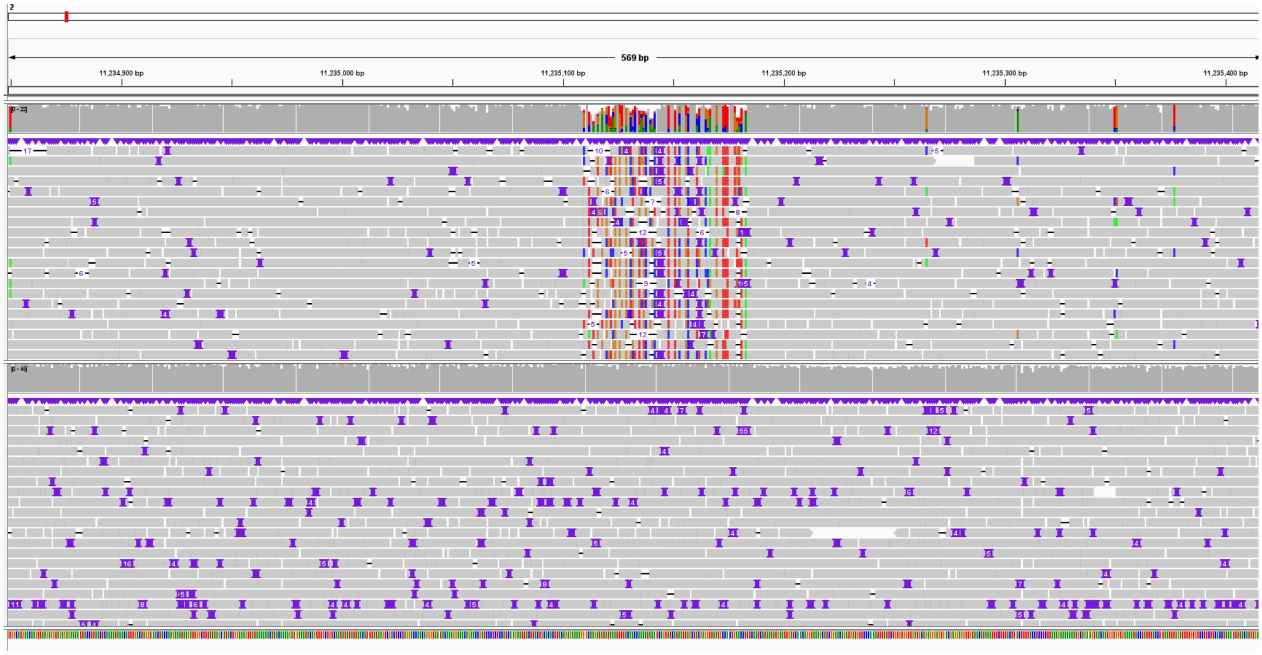

**Figure 22 |** CSUB chromosome 2, position 11235109, length 74 bp

Figure 23 & 24: This SV is a confirmed insertion, as can be seen in the overall alignment and the individual PacBio alignments.

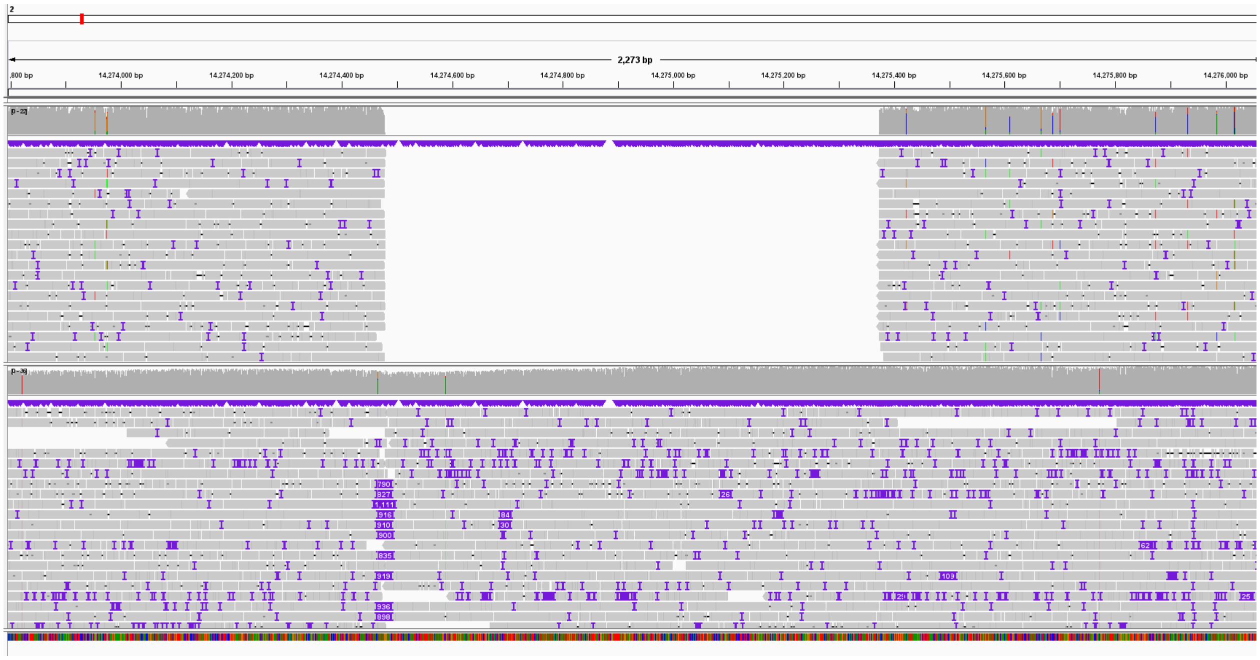

**Figure 23 |** CSUB chromosome 2, position 14274478, length 895 bp.

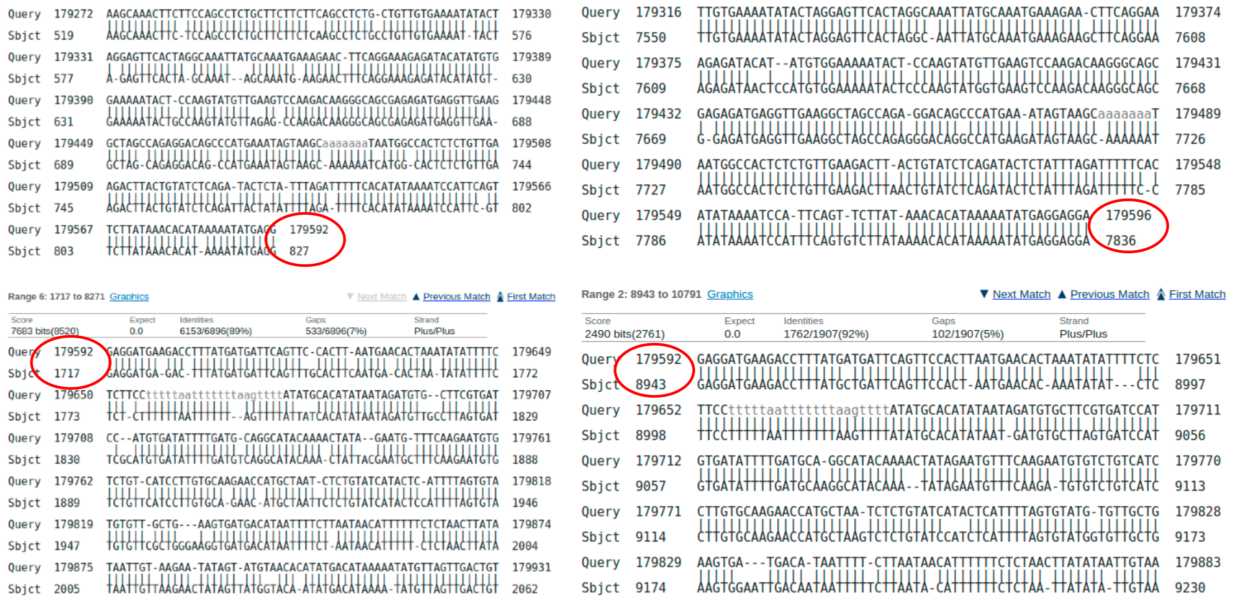


**Figure 24 |** Individual alignment of two PacBio reads for 'CSUB chromosome 2, position 14274478, length 895 bp'.

Figure 25 : We did not observe a SV at this position by inspecting the alignments.

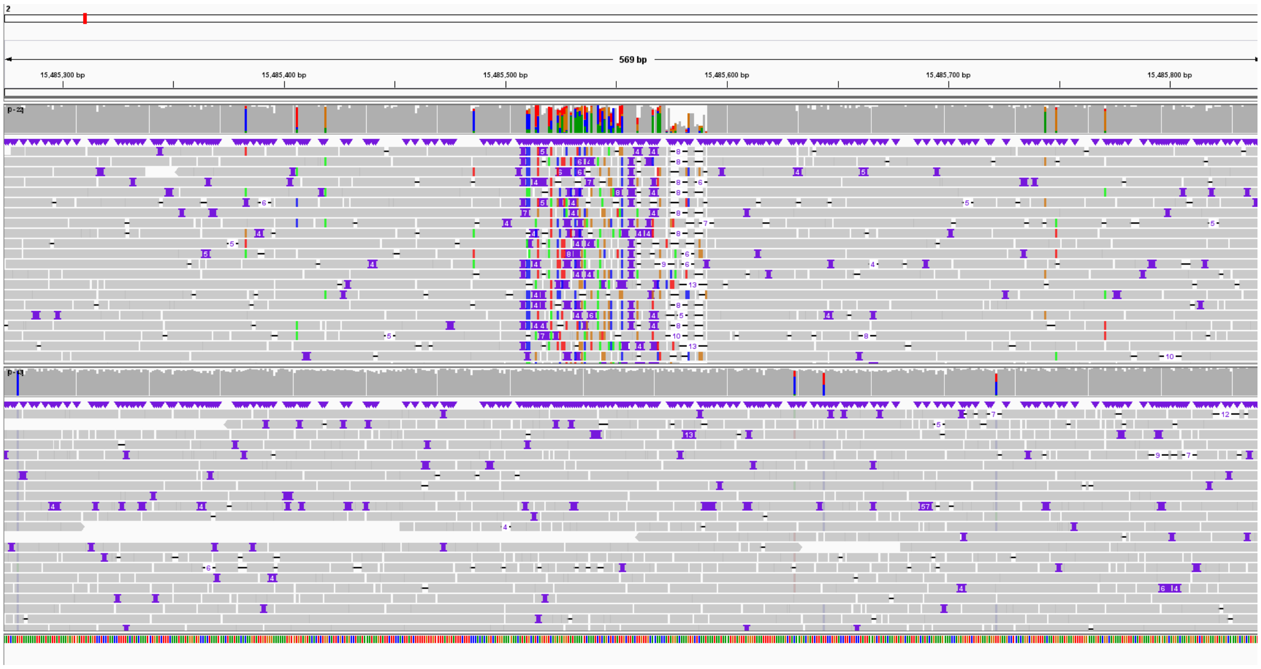


**Figure 25 |** CSUB chromosome 2, position 15485508, length 895 bp.

Figure 26 & 27: Insertions were observed across a tandem repeat region.

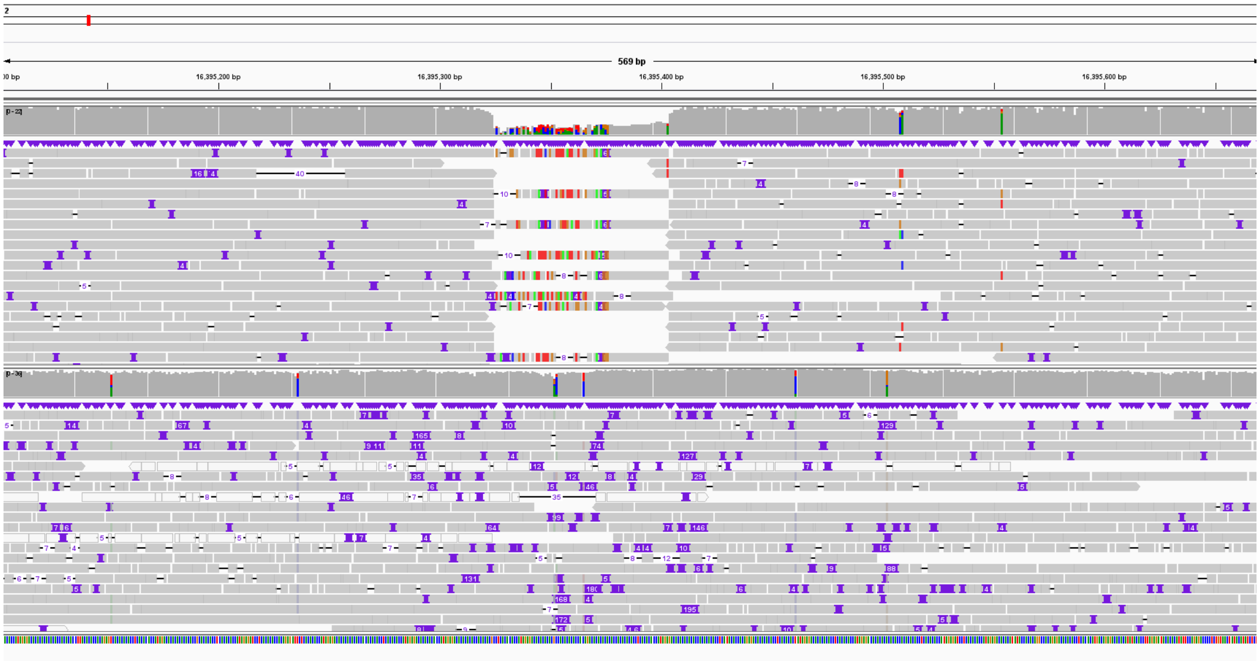

**Figure 26 |** CSUB chromosome 2, position 16395324, length 54 bp.

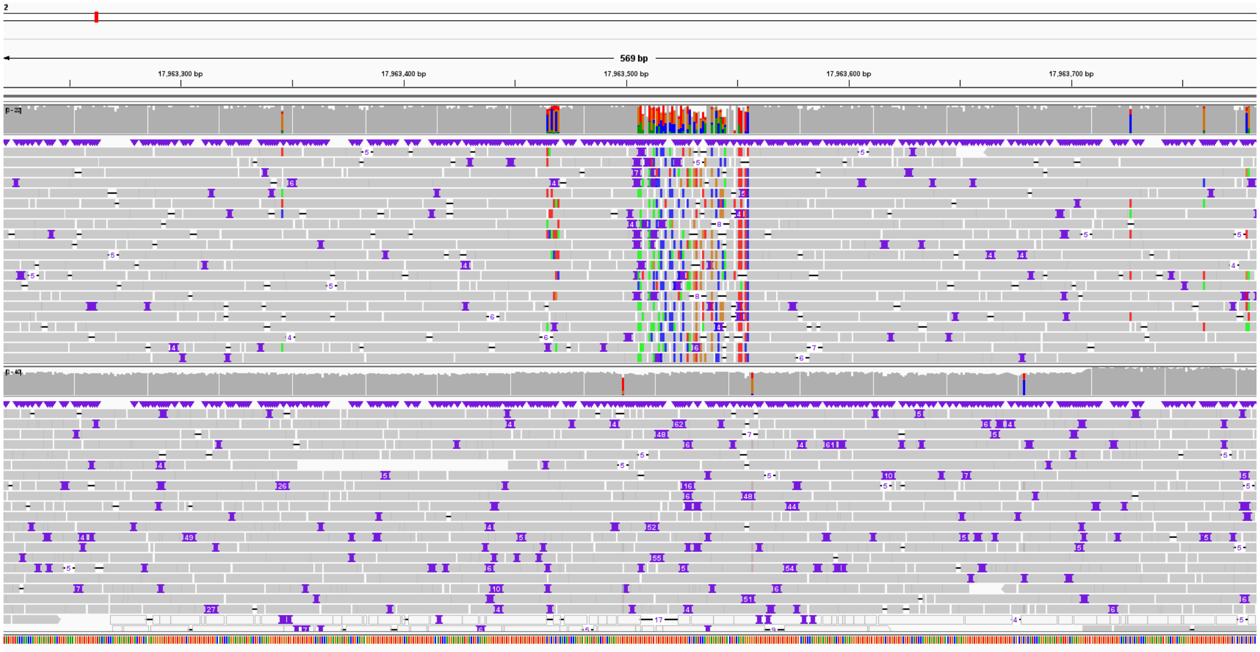

**Figure 27 |** CSUB chromosome 2, position 17963505, length 50 bp.

Figure 28: Insertions were observed across a repetitive region.

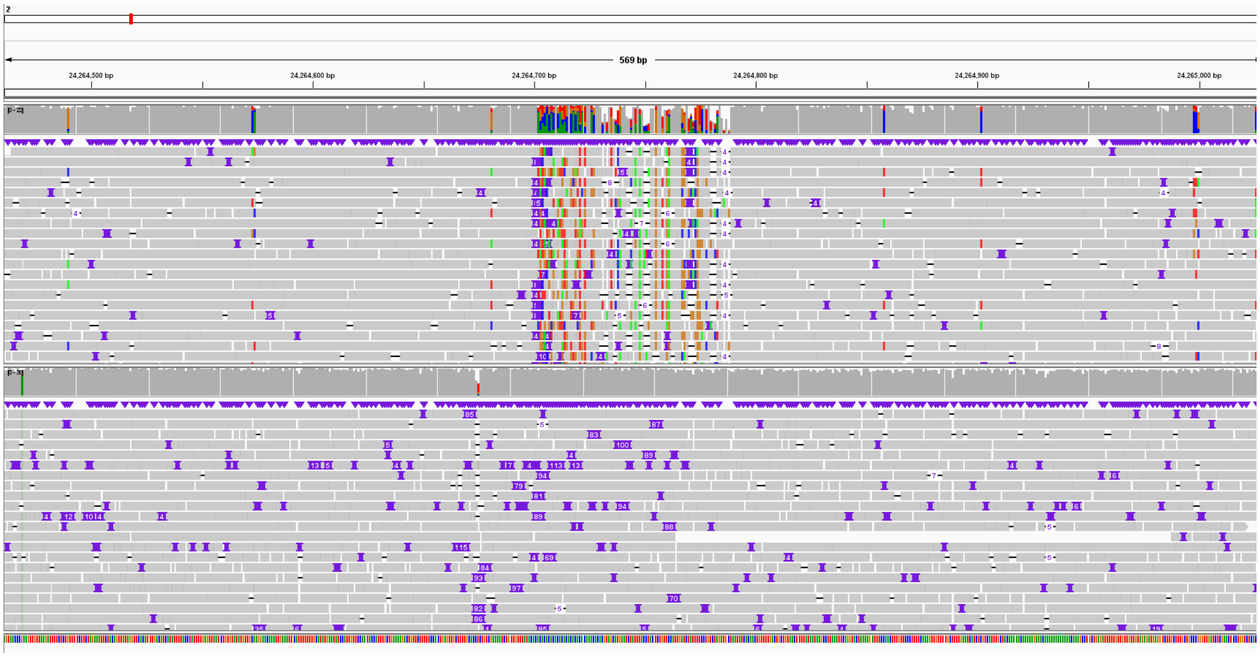


**Figure 28 |** CSUB chromosome 2, position 24264701, length 87 bp.

Figure 29 & 30 : We did not observe a SV at either position by inspecting the alignments.

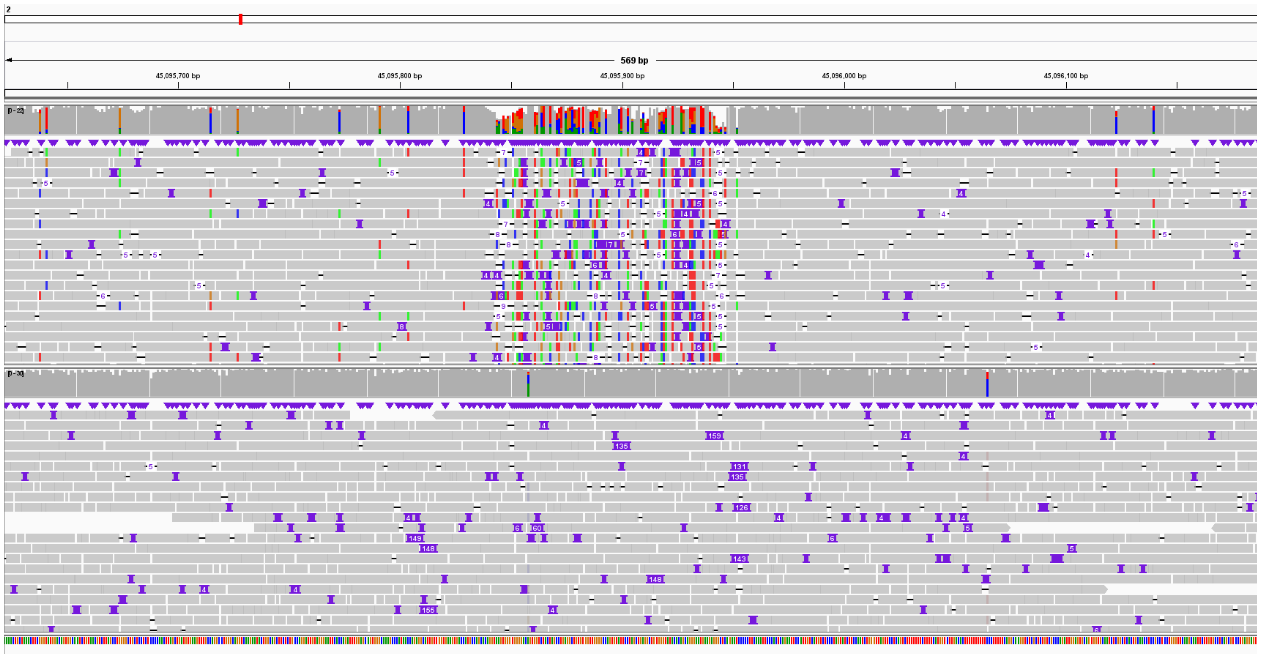


**Figure 29 |** CSUB chromosome 2, position 45095843, length 110 bp.

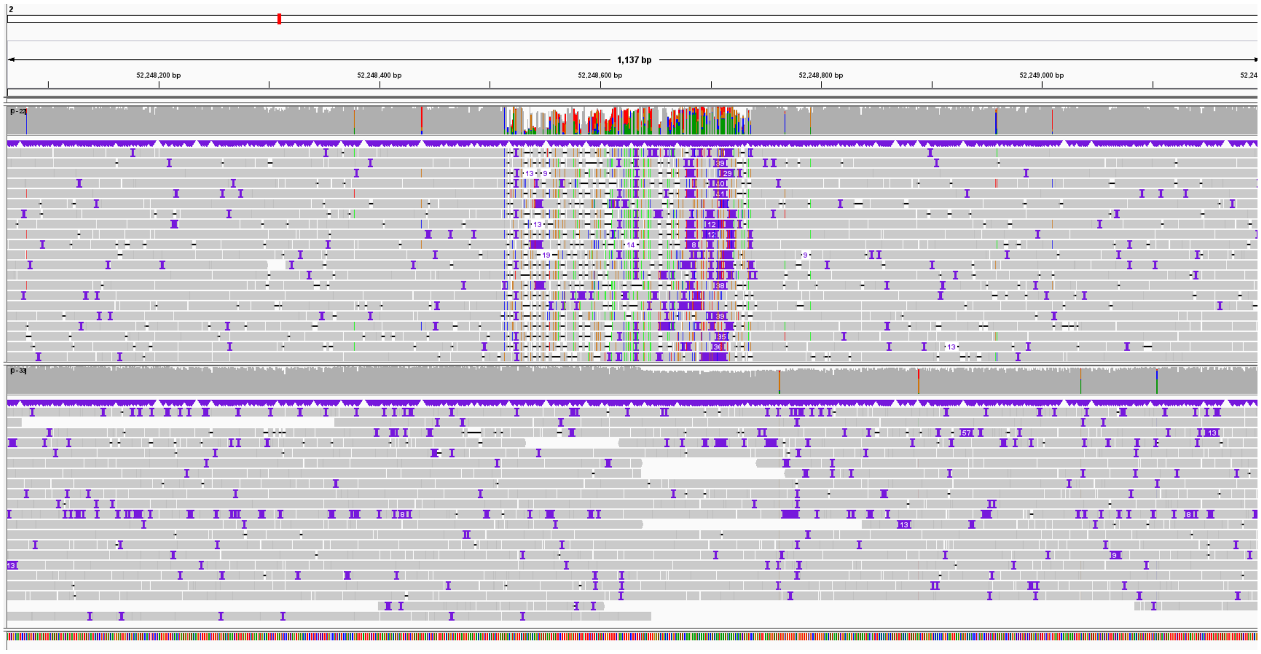

**Figure 30 |** CSUB chromosome 2, position 52248513, length 225 bp.

Figure 31 & 32: Insertions were observed across a repetitive region.

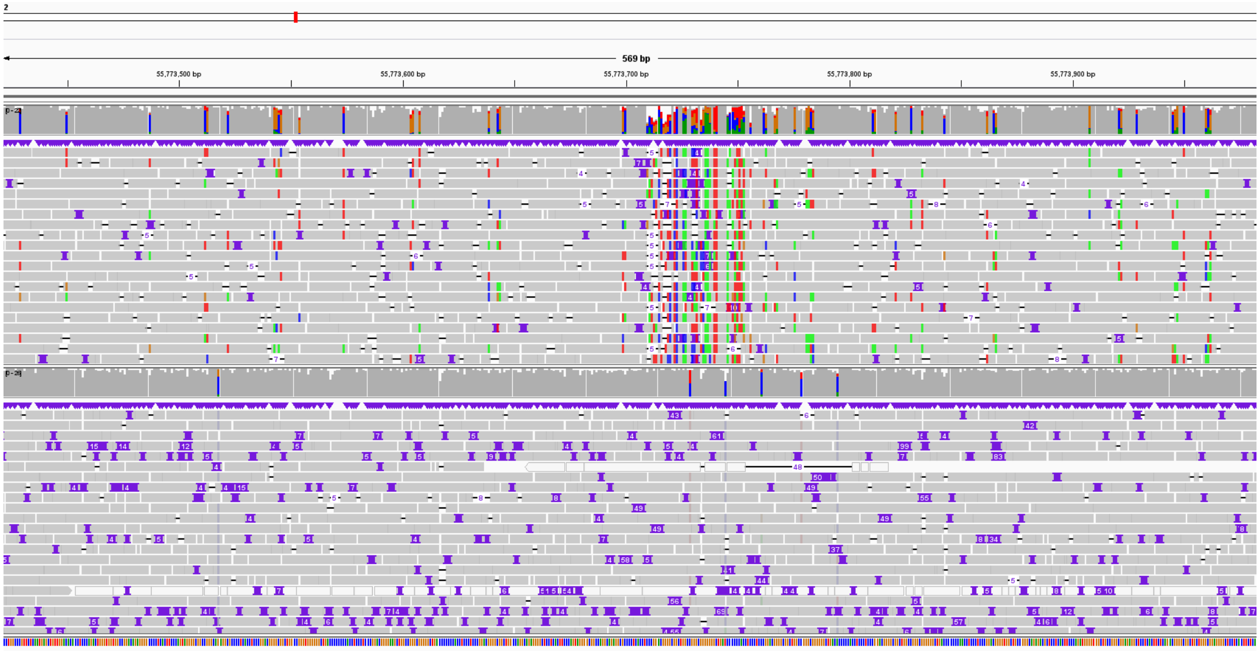

**Figure 31 |** CSUB chromosome 2, position 55773706, length 50 bp.

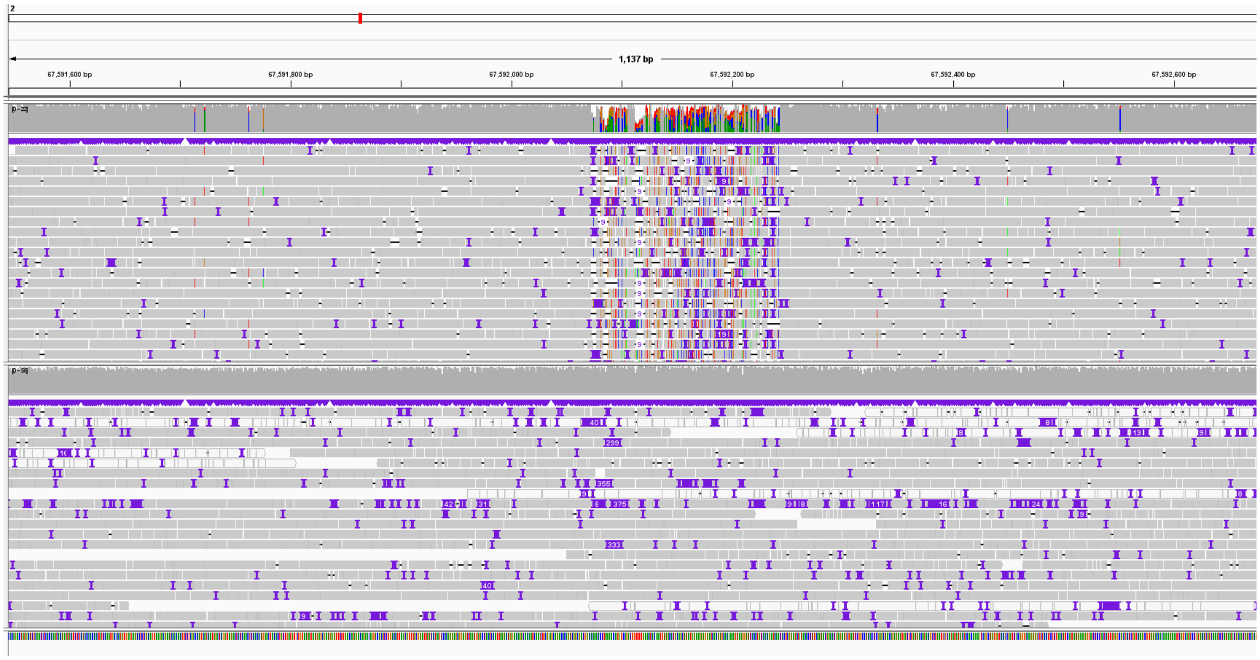


**Figure 32 |** CSUB chromosome 2, position 67592074, length 169 bp.

Figure 31: We observed an heterozygous deletion.

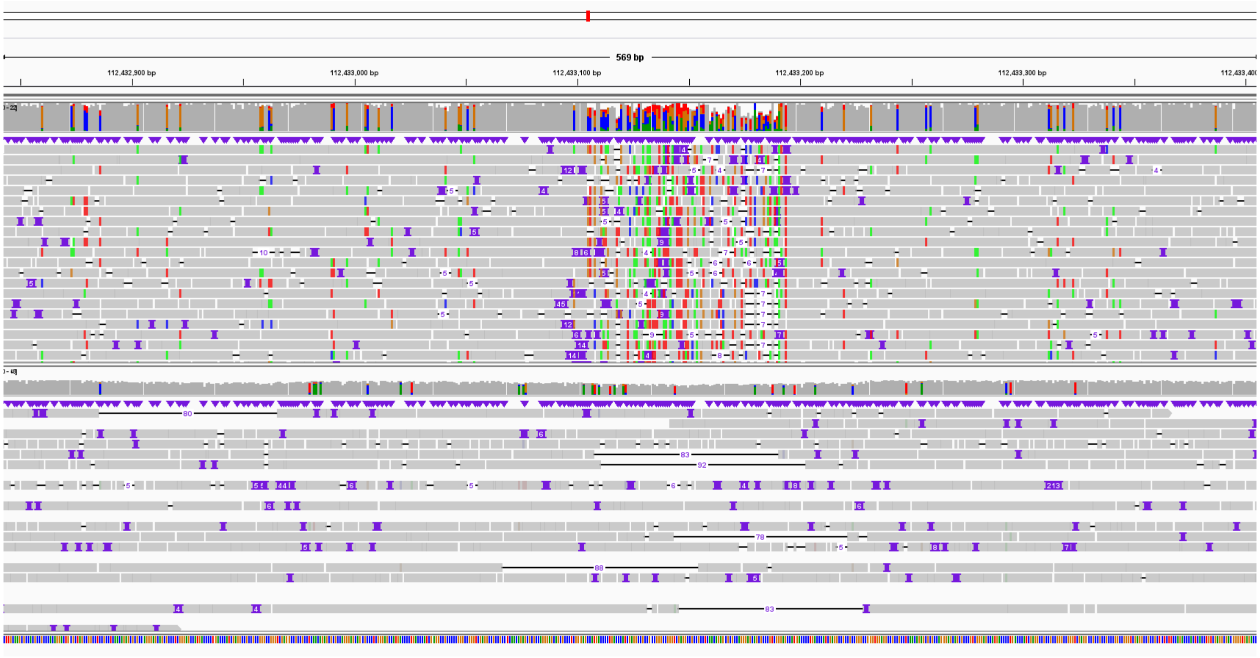


**Figure 31 |** CSUB chromosome 2, position 112433104, length 90 bp.

Figure 32: Insertions were observed across a tandem repeat region.

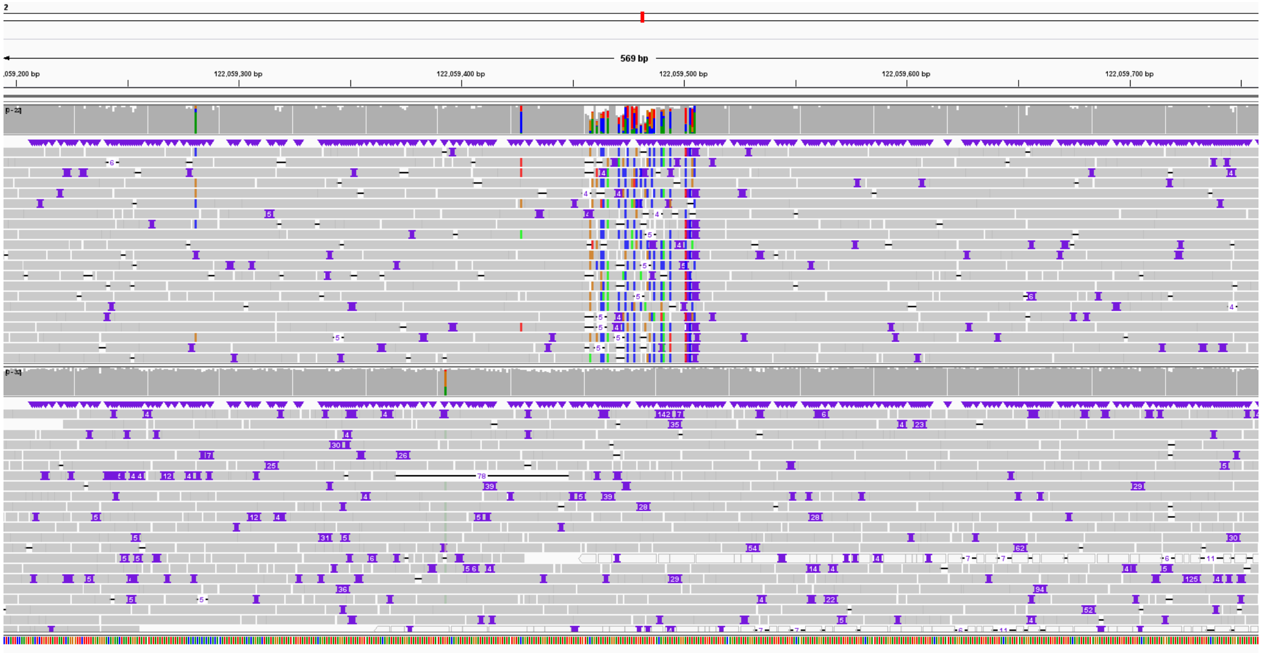


**Figure 32 |** CSUB chromosome 2, position 122059455, length 50 bp.
